# *parSMURF*, a High Performance Computing tool for the genome-wide detection of pathogenic variants

**DOI:** 10.1101/2020.03.18.994079

**Authors:** Alessandro Petrini, Marco Mesiti, Max Schubach, Marco Frasca, Daniel Danis, Matteo Re, Giuliano Grossi, Luca Cappelletti, Tiziana Castrignanò, Peter N. Robinson, Giorgio Valentini

**Affiliations:** AnacletoLab - Dipartimento di Informatica, Università degli Studi di Milano, Italy; Berlin Institute of Health (BIH), Berlin, Germany; Charité – Universitätsmedizin Berlin, Berlin, Germany; The Jackson Laboratory for Genomic Medicine, Farmington CT, USA; CINECA, SCAI SuperComputing Applications and Innovation Department, Roma, Italy

**Keywords:** High Performance Computing Tool for Genomic Medicine, Parallel Machine Learning Tool for Big Data, Parallel Machine Learning Tool for Imbalanced Data, Ensemble Methods, Machine Learning for Genomic Medicine, Machine Learning for Imbalanced Genomic Data, Prediction of Deleterious Variants, Prediction of Pathogenic Variants, High Performance Computing, Cluster of Computing Nodes, Mendelian Diseases, GWAS

## Abstract

Several prediction problems in Computational Biology and Genomic Medicine are characterized by both big data as well as a high imbalance between examples to be learned, whereby positive examples can represent a tiny minority with respect to negative examples. For instance, deleterious or pathogenic variants are overwhelmed by the sea of neutral variants in the non-coding regions of the genome: as a consequence the prediction of deleterious variants is a very challenging highly imbalanced classification problem, and classical prediction tools fail to detect the rare pathogenic examples among the huge amount of neutral variants or undergo severe restrictions in managing big genomic data. To overcome these limitations we propose *parSMURF*, a method that adopts a hyper-ensemble approach and oversampling and undersampling techniques to deal with imbalanced data, and parallel computational techniques to both manage big genomic data and significantly speed-up the computation. The synergy between Bayesian optimization techniques and the parallel nature of *parSMURF* enables efficient and user-friendly automatic tuning of the hyper-parameters of the algorithm, and allows specific learning problems in Genomic Medicine to be easily fit. Moreover, by using MPI parallel and machine learning ensemble techniques, *parSMURF* can manage big data by partitioning them across the nodes of a High Performance Computing cluster.

Results with synthetic data and with single nucleotide variants associated with Mendelian diseases and with GWAS hits in the non-coding regions of the human genome, involving millions of examples, show that *parSMURF* achieves state-of-the-art results and a speed-up of 80× with respect to the sequential version.

In conclusion *parSMURF* is a parallel machine learning tool that can be trained to learn different genomic problems, and its multiple levels of parallelization and its high scalability allow us to efficiently fit problems characterized by big and imbalanced genomic data.

**Availability and Implementation:** The C++ OpenMP multi-core version tailored to a single workstation and the C++ MPI/OpenMP hybrid multi-core and multi-node *parSMURF* version tailored to a High Performance Computing cluster are both available from github: https://github.com/AnacletoLAB/parSMURF

## Background

High throughput bio-technologies, and the development of Artificial Intelligence methods and techniques has opened up new research avenues in the context of the Genomic and Personalized Medicine [1, 2]. In particular Machine Learning [3], whole-genome sequencing (WGS) technologies [4, 5], and large population genome sequencing projects [6, 7] play a central role for the detection of rare and common variants associated with genetic diseases and cancer [8, 9].

In this context, while disease-associated variants falling in the protein-coding regions of the genome have been largely studied [10, 11, 12], this is not the case for disease-associated variants located in the non-coding regions of the genome, where our understanding of their impact on cis and trans-regulation is largely incomplete. Nevertheless, several studies ended up that most of the potential pathogenic variants lie in the non-coding regions of the human genome [13].

Driven by the aforementioned motivations many efforts have been devoted in recent years by the scientific community to develop reliable tools for the identification and prioritization of “relevant” non-coding genetic variants. CADD is one of the first machine learning-based method applied for this purpose on a genome-wide scale [14]. By combining different annotations into a single measure for each variant using firstly an ensemble of support vector machines and in the current version a fast and efficient logistic regression classifier, CADD likely represents the most used and well-known tool to predict deleterious variants [15].

Starting from this work other machine learning-based methods for the detection of deleterious or pathogenic variants have been proposed, ranging from multiple kernel learning techniques [16], to deep neural networks [17, 18], multiple kernel learning integrative approaches [16], unsupervised learning techniques to deal with the scarcity of available annotations [19], and linear models for functional genomic data combined with probabilistic models of molecular evolution [20]. Other approaches predicted the effect of regulatory variation directly from sequence using gkm-SVM [21] or deep learning techniques [22]. More details are covered in two recent reviews on machine learning methods for the prediction of disease risk in non-coding regions of the human genome [23, 24].

All these tools are faced with relevant challenges related to the rarity of non-coding pathogenic mutations. Indeed neutral variants largely outnumber the pathogenic ones. As a consequence the resulting classification problem is largely unbalanced toward the majority class and in this setting it is well-known that imbalance-unaware machine learning methods fail to detect examples of the minority class (i.e pathogenic variants) [25]. Recently several methods showed that by adopting imbalance-aware techniques we can significantly improve predictions of pathogenic variants in non-coding regions. The first one (GWAVA) applied a modified random forest [26], where its decision trees are trained on artificially balanced data, thus reducing the imbalance of the data [27]. A second one (NCBoost) used gradient tree boosting learning machines with partially balanced data, achieving very competitive results in the prioritization of pathogenic Mendelian variants, even if the comparison with the other state-of-the-art methods have been performed without retraining them, but using only their pre-computed scores [28]. The unbalancing issue has been fully addressed by ReMM [29] and *hyperSMURF* [30], through the application of subsampling techniques to the “negative” neutral variants, and oversampling algorithms to the set of “positive” pathogenic variants. Moreover a large coverage of the training data and an improvement of the accuracy and the diversity of the base learners is obtained through a partition of the training set and a hyper-ensemble approach, i.e. an ensemble of random forests which in turn are ensembles of decision trees. *hyperSMURF* achieved excellent results in the detection of pathogenic variants in the non-coding DNA, showing that imbalance-aware techniques play a central role to improve predictions of machine learning methods in this challenging task.

Nevertheless these imbalance-aware methods have been usually implemented with no or very limited use of parallel computation techniques, thus making problematic their application to the analysis of big genomic data. Furthermore, the *hyperSMURF* method is computationally intensive and characterized by a large number of learning parameters that need to be tuned to ensure optimal performance, thus requiring prohibitive computing costs, especially with big genomic data.

To address the aforementioned limitations, in this article we propose *parSMURF*, a novel parallel approach based on *hyperSMURF*. While other methods suitable for the assessment of the impact of variants located in protein-coding regions are able to run in parallel environments [31], this is not true for the assessment of non-coding variants. The main goal in the design and development of *parSMURF* is to make available to the scientific community a general and flexible tool for genomic prediction problems characterized by big and/or highly imbalanced data, while ensuring state-of-the-art prediction performance. The high computational burden resulting by the proper tuning and selection of the learning (hyper)-parameters is addressed through a scalable and parallel learning algorithm that leverages different levels of parallelization, and a Bayesian optimizer for their automatic and efficient tuning.

In the remainder we present the *parSMURF* algorithm, its relationships with its sequential version *hyperSMURF*, and its two different implementations respectively for multi-core workstations and for a highly parallel High Performance Computing cluster. In the Results section experiments with big synthetic and actual genomic data show that *parSMURF* scales nicely with big data and significantly improves the speed-up of the computation. Finally experiments with Mendelian data and GWAS hits at whole-genome level show that *parSMURF* significantly improves over its sequential couterpart *hyperSMURF*, by exploiting its multiple levels of parallelism and the automatic tuning of its learning hyper-parameters through a grid search or a Bayesian optimization method. *parSMURF*^1^ multi-thread and hybrid multi-thread and multi-process MPI C++ *parSMURF*^*n*^ implementations are well-documented and freely available for academic and research purposes.

## Methods

*Parallel SMote Undersampled Random Forest* (*parSMURF*) is a fast, parallel and highly scalable algorithm designed to detect deleterious and pathogenic variants in the human genome. The method is able to automatically tune its learning parameters even with large data sets, and to nicely scale with big data.

Starting from the presentation of the characteristics and limitations of *hyperSMURF* [30], in this section we introduce the parallel algorithm *parSMURF* and its two variants named *multi-core* parSMURF (*parSMURF*^1^) and *multi-node* parSMURF (*parSMURF*^*n*^). The first one is suitable for the execution on a single workstation that features one or more multi-core processors, while the second one is designed for a High Performance Computing cluster, where the computation is distributed across several interconnected nodes. Although developed for different hardware architectures, they both share the same core parallelization concepts and the same chain of operations performed on each parallel component of the algorithm. Finally, we discuss the computational algorithms proposed to automatically learn and tune the *parSMURF* hyper-parameters in order to properly fit the model to the analyzed genomic data.

### From *hyperSMURF* to *parSMURF*

*hyperSMURF* is a supervised machine learning algorithm specifically designed to detect deleterious variants where variants associated with diseases are several order of magnitude lesser than the neutral genetic variations. *hyperSMURF* tackles the imbalance of the data using three learning strategies:

- balancing of the training data by oversampling the minority class and undersampling the majority class;
- improving data coverage through ensembling techniques;
- enhancing the diversity and accuracy of the base learners through hyper-ensembling techniques.

The high-level logical steps of the *hyperSMURF* algorithm are summarized in Figure 1. At step (1) *hyperSMURF* creates several sets of training data by using all the available examples of the minority (positive) class and partitioning the set of the majority (negative) class: as a result each set includes all the positive examples and a subset of the majority (negative) class. From this point on, each training set is processed independently. In step (2), examples of the minority class are over-sampled through the SMOTE algorithm [32] while examples of the majority class are undersampled according to an uniform distribution. Each training set is is now formed by a comparable number of positive and negative examples and it can be used in step (3) to train the random forest. This process is applied to all the parts of the partition of the original training set, thus generating an ensemble of random forest models. At step (4) all the predictions separately computed by each trained model are finally combined to obtain the “consensus” prediction of the hyper-ensemble.

**Figure 1.**
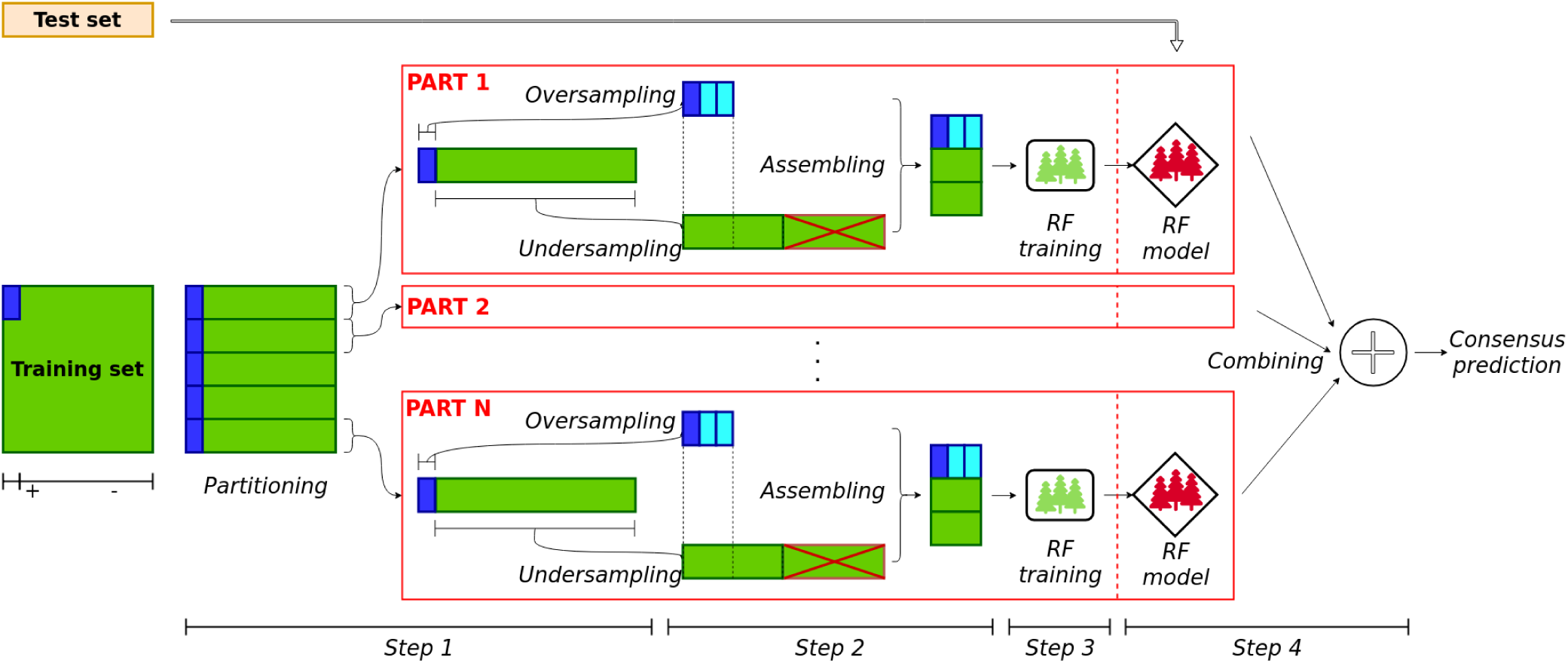
High-level scheme of *hyperSMURF*. Step (1): partitioning of the training set (the minority/positive class is represented in blue, while the majority/negative class is in green). Step (2): application of oversampling and undersampling approaches, and assembling of the training set. Step (3): training of the RF models. Step (4): Testing and aggregation of predictions outcomes.

The behavior of the algorithm strongly depends on the learning hyper-parameters, reported in Table 1, which deeply influence the *hyperSMURF* performance, as shown in [33], and fine tuning of the learning parameters can dramatically improve prediction performance. Since *hyperSMURF* is an ensemble of random forests which in turn are ensembles of decision trees, its sequential implementation undergoes a high execution time, especially on large datasets, thus limiting a broad exploration of the hyper-parameter space. Moreover *hyperSMURF* cannot be easily applied to big data, due to its time and space complexity issues.

**Table 1.** *hyperSMURF* learning hyper-parameters

| Parameters | Description |
| --- | --- |
| nParts | Number of parts of the partition |
| fp | Multiplicative factor for oversampling the minority class. For instance with $fp = 2$ two novel examples are synthesized for each positive example of the original data set, according to the SMOTE algorithm. |
| ratio | Ratio for the undersampling of the majority class. For instance, $ratio = 2$ sets the number of negatives as twice the total number of original and oversampled positive examples. |
| k | Number of the nearest neighbors of the SMOTE algorithm |
| nTrees | Number of trees included in each random forest |
| mtry | Number of features to be randomly selected at each node of the decision trees included in the random forest |

Nevertheless, looking at Figure 1, we can observe that *hyperSMURF* is characterized by the following features:

i. the same operations (over- and under-sampling, data merging, training and model generation and prediction evaluation) are performed over different data belonging to different partitions;
ii. the operations performed over different data are independent, i.e. there is no interaction between the computation of two different partitions;
iii. the algorithm does not require any explicit synchronization during the elaboration of two or more partitions.

Putting together these observations, we can redesign *hyperSMURF* leveraging its intrinsic parallel nature and using state-of-the-art parallel computation techniques. The resulting newly proposed *parSMURF* algorithm is schematically summarized in Algorithm 1. The parallelization is performed by grouping parts of the partition in *chunks* (see also Figure 2). The *parSMURF* parameter *q* (number of chunks) determines at high level the parallelization of the algorithm, i.e. how many chunks can be processed in parallel.

**Figure 2.**
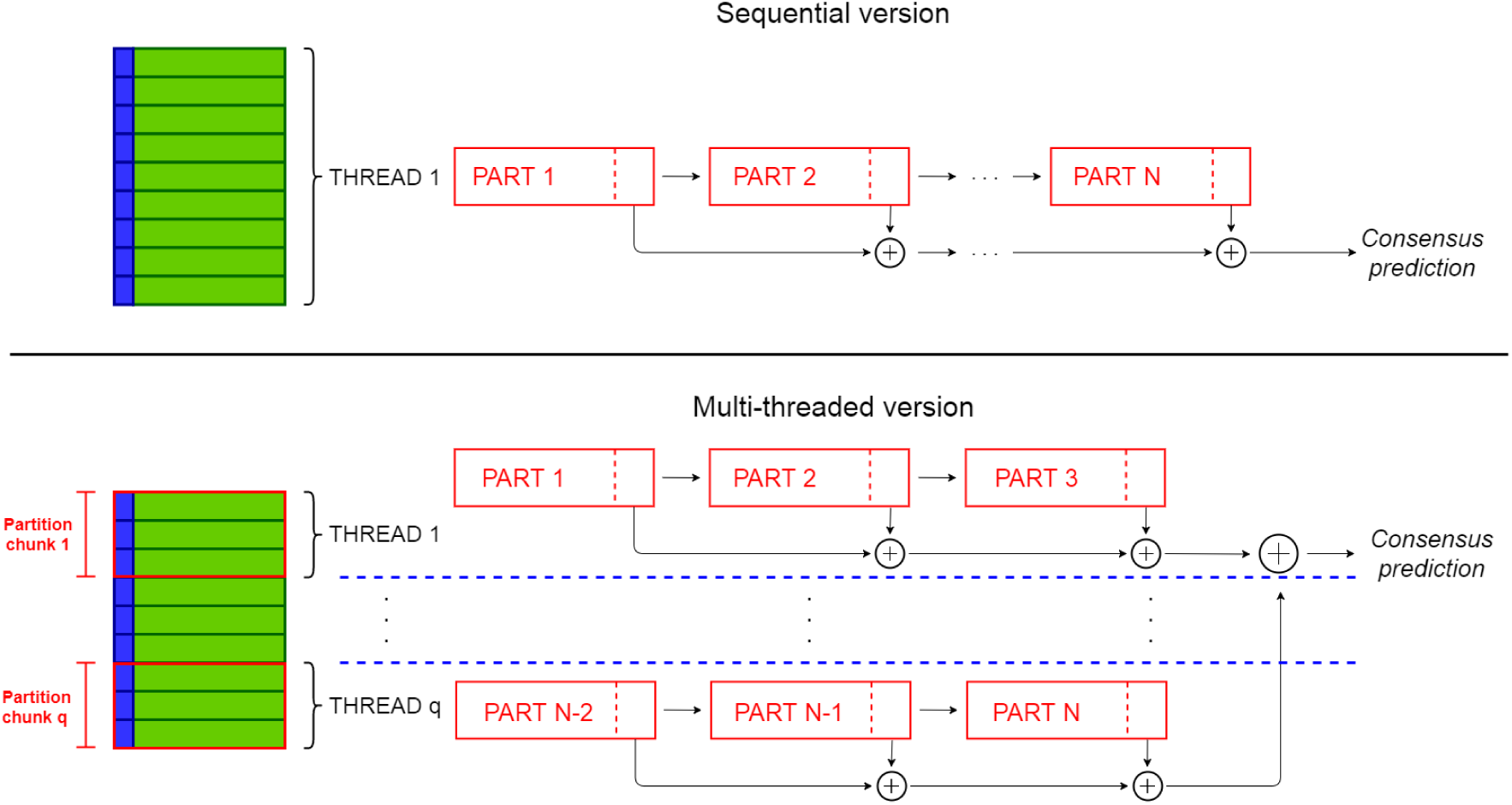
Comparison between the sequential *hyperSMURF* (top) and multi-core *parSMURF*^1^ (bottom) execution schemes.

During training, the main activities of the *parSMURF* algorithm are executed in parallel for each chunk (outer `for` loop in Algorithm 1). A further level of parallelism can be realized through the inner `for` loop where each part 𝒩 ^*′*^ of the chunk 𝒞_*i*_ undergoes a parallel execution. Note however that “parallel” in the inner `for` loop is in brackets to highlight that this second level of parallelization can or cannot be implemented, according to the complexity of the problem and the available underlying computational architecture.

#### Algorithm 1 *parSMURF* algorithm (training)

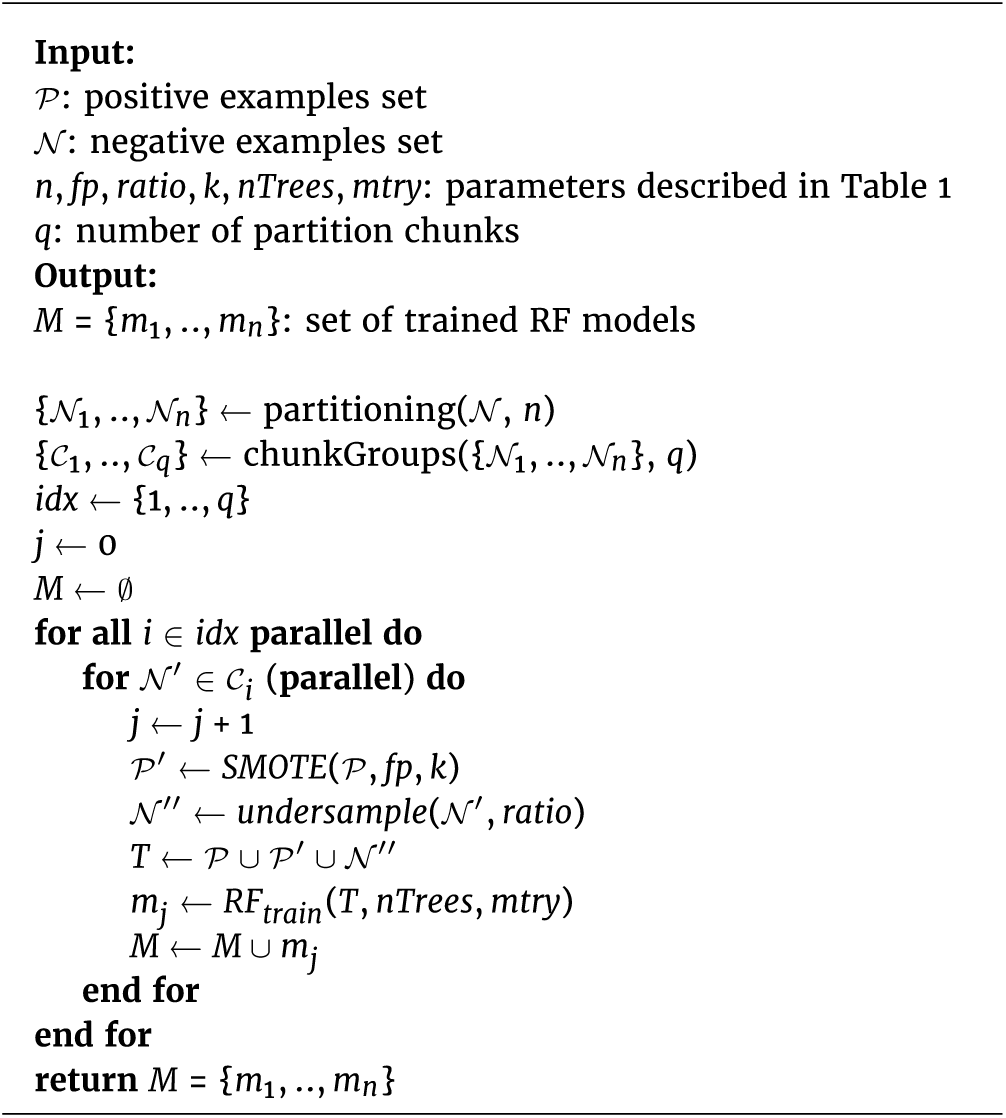

Algorithm 2 also shows that hyper-ensemble predictions conducted during testing can be easily performed through parallel computation: each model can be tested independently over the same test set and the consensus prediction is computed by averaging the ensemble output.

#### Algorithm 2 *parSMURF* algorithm (testing)

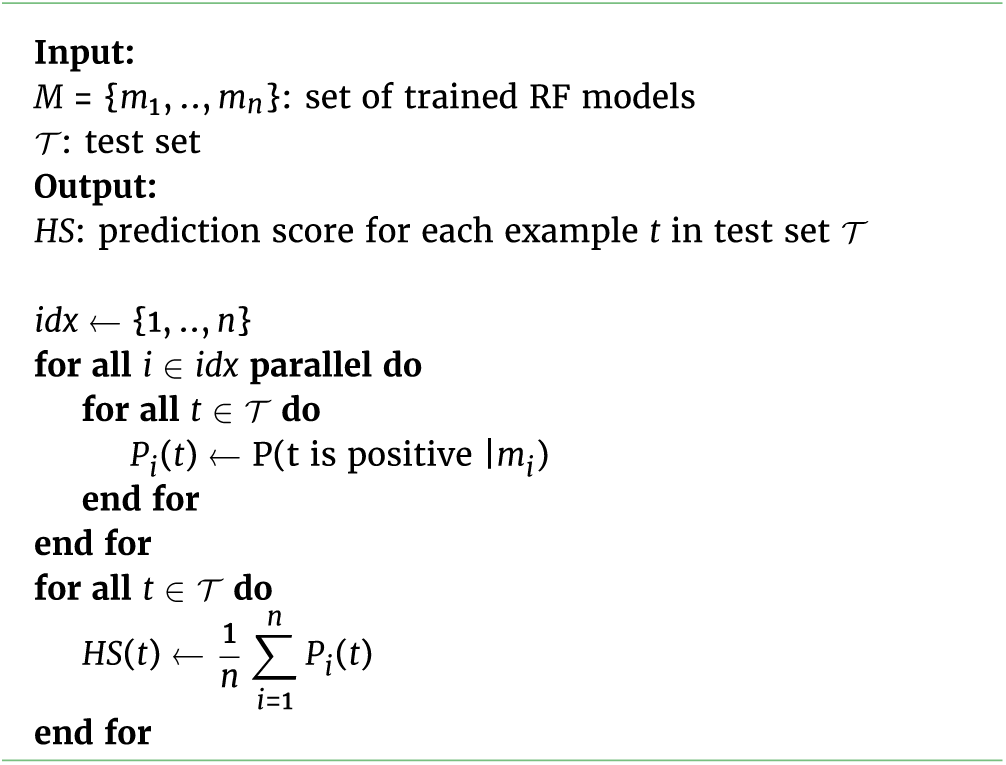

### Multi-core *parSMURF*^1^

The idea on which multi-core *parSMURF* builds upon is that all operations performed on the different parts of the partition can be assigned to multiple core/threads and processed in parallel. Namely, given *q* threads, the data parts *N*1, …, *N*_*n*_ are equally distributed among threads so that thread *i* receives a subset (chunk) *C*_*i*_ of parts, and processes its assigned parts in sequence. Since each partition chunk is assigned to its own thread, chunk processing is performed in parallel with architectures featuring multiple processing cores.

In Figure 2 (top) a scheme of the execution of the sequential *hyperSMURF* is shown: each partition is processed sequentially and the output predictions are accumulated as the computation goes on. On the contrary, with *parSMURF*^1^ (Figure 2, bottom), chunks of partitions are assigned to different execution threads and are processed in parallel. To avoid data races, each thread accumulates partial results, and then the master thread collects them once the computation of each thread is ended. Moreover, each thread keeps only a local copy of the subset of the data which is strictly required for its computation; this minimizes memory consumption and, at the same time, does not impose any need for synchronization between concurrent threads.

This scheme is expected to show a remarkable speedup with the increase of processing cores and the available local memory of the system. Since parallelization occurs at “partition chunk” level, instances of *parSMURF*^1^ with a reduced partition size benefits only partially of a multicore execution. On the other side, partitions characterized by a very high number of parts can theoretically scale well with the number of processing cores but, unfortunately, current processors have constraints in the number of available cores. Moreover, big data computation may exceed the storage capacity of a single workstation, thus making the application of *parSMURF*^1^ in this experimental context problematic.

### Multi-node *parSMURF*^*n*^

This version of *parSMURF* has been designed to process big data, to both improve speed-up and make feasible computations that exceed the storage capacity of a single workstation. Moreover *parSMURF*^*n*^ allows the fine tuning of the model parameters even when big data should be analyzed.

#### Architecture

As for the multi-core version, *parSMURF*^*n*^ exploits parallelization at partition level, but also introduces a second level of parallelization: the higher level is performed through the computing nodes of a cluster, i.e. a set of computing machines inter-connected by a very fast networking infrastructure; the lower level is realized through multi-threading at single node level by exploiting the multi-core architecture of each single node of the cluster. In this novel scheme, each node receives a partition chunk, which is processed in parallel with the other chunks assigned to the remaining nodes. Chunks in turn are further partitioned in sub-chunks, distributed among the computing cores available at the local node. Finally an optional third level of parallelization is available by assigning multiple threads to the random forests which process the different parts of the partition (recall that a random forest is in turn an ensemble of decision trees).

The higher level of parallelization leverages the MPI programming paradigm and standard [34] to transfer information among nodes. This programming paradigm requires that several instances of the same program are executed concurrently as different processes (*MPI processes*) on different nodes interconnected by a network. Being different instances of the same program, each MPI process has its own memory space, therefore intercommunication, i.e. data exchange between processes, occurs explicitly by invoking the corresponding actions, as required by the MPI standard.

*parSMURF*^*n*^ adopts a master–slave setting, with a master process coordinating a set of MPI slave processes, also called *worker processes*, which in turn manage the partition computation. Master and worker roles are described below:

- the *master process* is responsible for processing the command line parameters, loading data in its memory space, generating partition and chunks, sending the proper subset of data to each worker process (including the test set and the proper fraction of the training set) and finally collecting and averaging their output predictions.
- each *worker process* realizes the computation on the assigned chunk of partitions, generates sub–chunks of its own chunk and processes them through multi-threading - i.e. distributes the computation of the sub–chunks over the available computing threads - and sends the output predictions back to the master process.

We point out that in principle *parSMURF*^*n*^ can be executed also on a single machine, where multiple copies of the same program are processed by the available cores, but in this case it undergoes the same limitations of *parSMURF*^1^. Figure 3 provides a high level scheme of the execution of *parSMURF*^*n*^.

#### parSMURF^*n*^ *intercommunication*

Figure 4 shows a schematic view of the intercommunication between *parSMURF* MPI processes.

**Figure 3.**
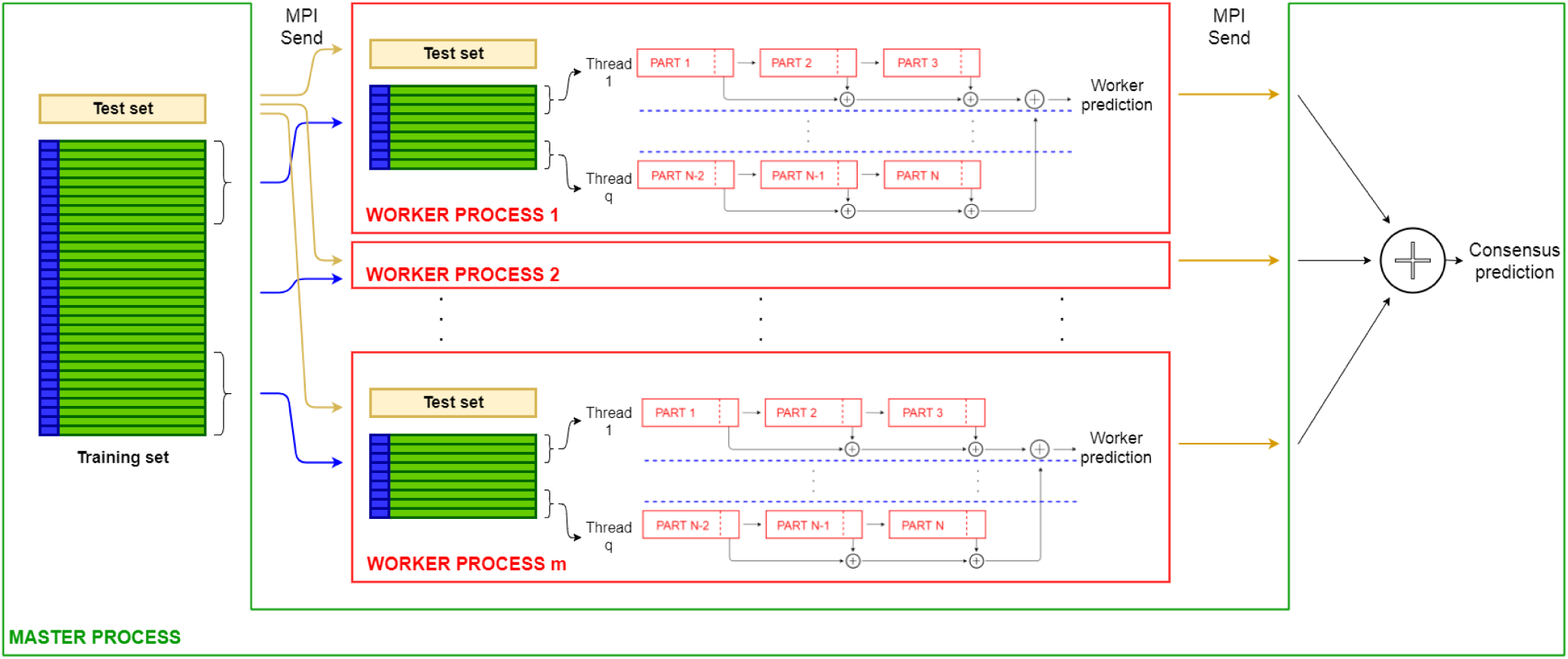
High-level scheme of the multi-node *parSMURF*^*n*^ implementation.

**Figure 4.**
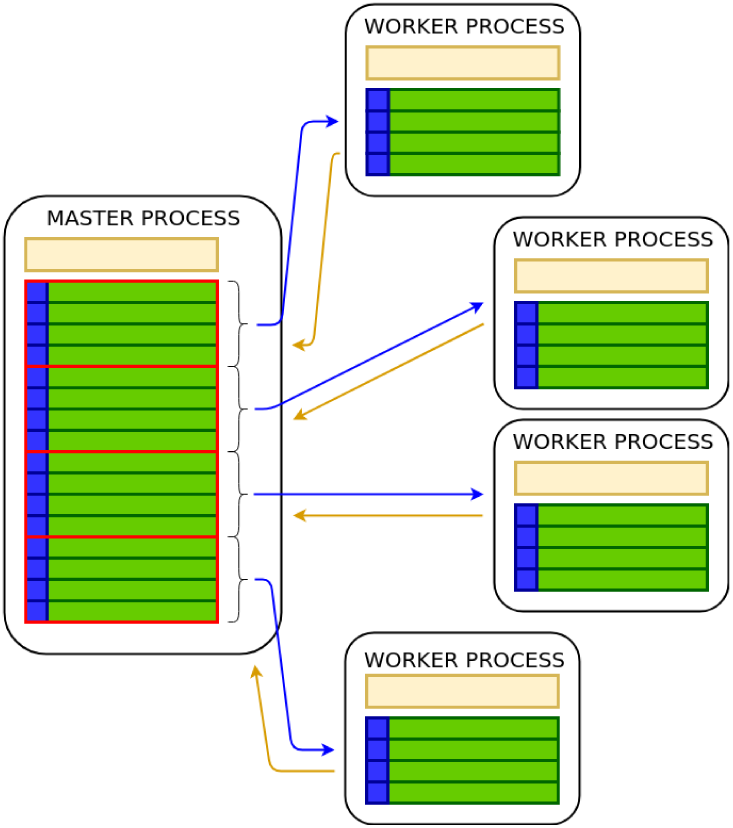
High-level intercommunication scheme between MPI processes in multi-node *parSMURF*^*n*^. Blue arrows represent data flows from the master process to worker processes (different chunks of partitions and the same test set) and yellow arrows represent data flows from the worker processes to the master (output predictions).

The computation in the worker processes is performed as in the multi-core version of *parSMURF*, except for a slight difference in the subsampling of the majority class, since this operation is no longer executed by the worker processes but by the master instead. Indeed, by observing that subsampling requires some examples to be discarded, there is no need of sending to the worker processes an entire part of the partition, but only the selected subset of examples. This design choice minimizes the amount of data to be sent to a worker process, since for each partition only the positive samples (that are going to be oversampled in the worker process) and the already under-sampled negative examples are sent to the worker processes.

In an ideal parallel application, computing nodes should never interact, since every data exchange creates latencies that affect the overall *occupancy* - i.e. the ratio between the amount of time a computing node is processing data and the total execution time. However, in real applications this rarely happens, and data have to be exchanged between processes. As a general rule, communication should be minimal in number and maximal in size, since the following equality holds:

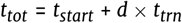

where *t*_*tot*_ is the total time for the data send, *d* is the amount of data in bytes to be transferred, *t*_*trn*_ is the time required to transfer one byte of data and *t*_*start*_ is the time required by the interconnecting networking system for beginning a communication between nodes. *t*_*start*_ is constant, therefore transferring data as a big chunk is generally faster than several smaller ones, since *t*_*start*_ penalty is paid only once in the former case. However, in real world MPI parallel applications, the main objective is to parallelize computation to speed-up execution, and maximum efficiency is achieved by overlapping data transmission and computation. This is easier to obtain when data is *streamed*, i.e. sent in small chunks which are consumed as soon as they reach the receiver MPI process: in this way we can minimize the inactivity time of a node, waiting for data to be received.

#### *Maximizing* parSMURF^*n*^ *performance*

For maximizing performances *parSMURF*^*n*^ adopts the following strategies to find the optimal balance between the size and number of data transmissions:

- *maximize occupancy*,
- *reduce the number of data send* or *broadcast*,
- *minimize latency*.

To *maximize occupancy*, the master process does not send the entire chunk of partitions to each worker process in a big data send; instead, parts are sent one by one. This choice is ideal in the context of multithreading in worker processes: supposedly, given a partition with *n* parts and a number *x* of computing threads assigned to a working process, the master at first sends to each worker process *x* parts of its assigned chunk. When a worker thread finishes the computation of a part, another one is sent by the master for processing. This process goes on until the chunk is exhausted.

To *reduce the number of data send* or *broadcast* - i.e. one MPI process sending the same data to all the other processes - for each part, the master process assembles an array having all the relevant data required for the computation, i.e. the positive and negative examples (already subsampled, as stated earlier) and the parameters needed for the computation. Hence with just one MPI send operation, a part of the partition with its parameters is transferred to the worker process. Also, partial results of the predictions are locally accumulated inside each worker and sent to the master once the jobs for the assigned chunk are finished.

To *minimize latency*, the assembly of the data to be sent is multithreaded in the master process. In instances characterized by relatively small datasets and a high number of parts in the partition, it may happen that the master could not prepare and send data fastly enough to keep all the worker processes busy. For instance, a thread in a worker process may finish the computation for a part before data for the next one arrive, leaving the thread or the entire process inactive. This has been solved by spawning a number of threads in the master process equal to the number of worker processes the user has requested, each one assigned for preparing and sending the data to the corresponding worker. However, since memory usage in the master process can be greatly affected, an option is provided for disabling multithreading in the master. In this case, only one thread takes care of this task and parts are sent in round robin fashion to the working processes: this has been experimentally proven to be effective for those instances that require a particularly high memory usage.

### Hyper-parameters tuning

As in most ML methods, the accuracy of the predictions of the *parSMURF* models are directly related to the set *S* of hyper-parameters that control its learning behaviour. Hence, to maximize the usefulness of the learning approach, the value of each hyper-parameter of the set *S* must be chosen appropriately. In *parSMURF* the hyper-parameter set is composed by the 6-tuple of parameters reported in Table 1. Each parameter is discretized and constrained between a maximum and minimum value, hence the hyper-parameter space ℋ is a discrete six-dimensional hypercube. The validation procedure for the evaluation of each *h* ∈ ℋ is the internal cross-validation process, and the objective function (performance metrics) which has to be maximized is the Area Under the Precision Recall Curve (AUPRC).

*parSMURF* features two modes for automatically finding the hyper-parameters combination *h*_0_ ∈ ℋ that maximizes the model accuracy (parameter auto-tuning). The first strategy is a grid search over *H*, where each *h* ∈ ℋ is evaluated by internal cross-validation. This strategy is generally capable of finding values close to the best hyper-parameters combination, but it is very computationally intensive and suffers from the curse of dimensionality. The second strategy is based on a Bayesian Optimizer (BO) which iteratively builds a probabilistic model of the objective function *f* : ℋ → ℝ^+^ (in *parSMURF*, the AUPRC) by evaluating a promising hyper-parameter combination at each iteration and stopping when a global maximum of *f* is obtained. This procedure is less computationally intensive than the grid search and may also outperform the latter in terms of prediction effectiveness. The Bayesian optimizer is based on [35] and its implementation “Spearmint-lite” [36] is included in the *parSMURF* package.

The whole procedure is summarized in Algorithm 3. Briefly, *parSMURF* provides the automatic tuning of the hyper-parameters in a context of internal *n*-fold cross-validation, and the Bayesian Optimizer (BO) is invoked in the while loop.

#### Algorithm 3 Automatic hyper-parameters tuning in *parSMURF* featuring Bayesian Optimization

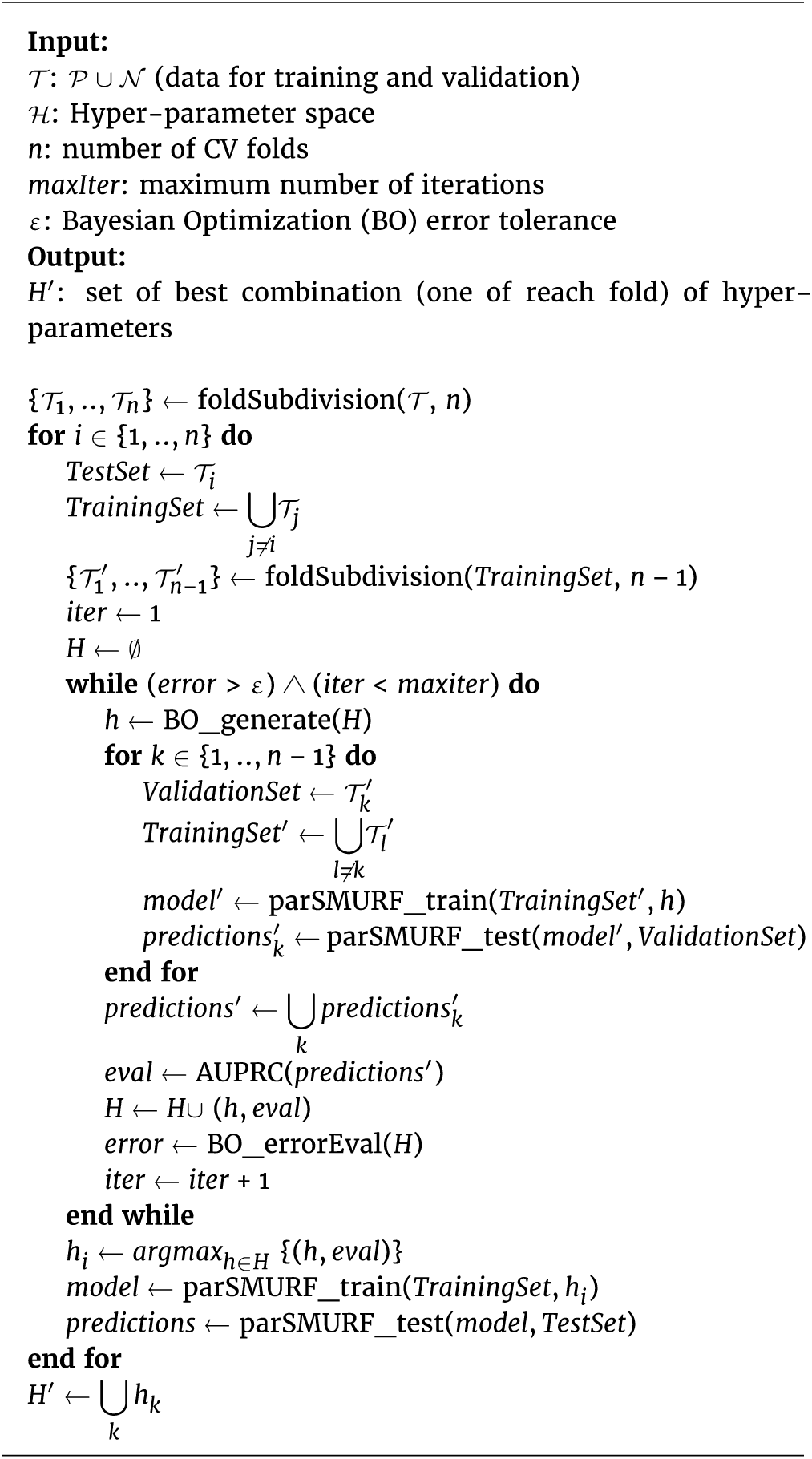

At each iteration, a new hyper-parameter combination *h* ∈ ℋ is generated by taking into account all the previously evaluated *h*. A new model is then trained and tested in the internal cross-validation procedure by using the newly generated *h*. The quality of the prediction is evaluated by means of AUPRC, and the tuple (*h, eval*) is submitted back to the Bayesian Optimizer for the generation of the next *h*. The while loop ends when the Bayesian Optimizer finds a probable global maximum (no further improvement in the error evaluation) or when the maximum number of iterations is reached. Grid search works in a similar way, but all *h* ∈ ℋ are exhaustively tested in the internal cross-validation phase.

## Results and Discussion

We applied *parSMURF* to both synthetic and real genomic data to show that the proposed method is able to:

- scale nicely with big data;
- auto-tune its learning parameters to optimally fit the prediction problem under study;
- improve *hyperSMURF* as well as other state-of-the-art methods in the prediction of pathogenic variants in Mendelian diseases and of regulatory GWAS hits in the GWAS Catalog.

All the experiments have been performed on the Cineca Marconi Supercomputing system [37], specifically using its Lenovo NeXtScale architechture, with 720 nodes, each one having 128 GBytes of RAM and 2 × 18-cores Intel Xeon E5-2697 v4 (Broad-well) CPUs at 2.30 GHz. The interconnecting infrastructure is a Intel Omnipath featuring 100 Gb/s of networking speed and a fat-tree 2 : 1 topology.

### Datasets

Genomic data are highly imbalanced toward the majority class, since the SNVs annotated as pathogenic represent a tiny minority of the overall genetic variation. Synthetic data have also been generated to obtain a high imbalance between positive and negative examples, in order to simulate the imbalance that characterizes several types of genomic data.

Synthetic data have been randomly generated using a spheric gaussian distribution for each of the 30 features. Among them only 4 are informative in the sense that the means of positive and negative examples are different, while all the other features share the same mean and standard deviation with both positive and negative examples. Synthetic data, as well as the R code for their generation are available from the GITHUB repository [38].

As an example of application of *parSMURF* to real genomic data, we used the dataset constructed in [29] to detect Single Nucleotide Variants (SNVs) in regulatory non-coding regions of the human genome associated with Mendelian diseases. The 406 positive examples are manually curated and include mutations located in genomic regulatory regions such as promoters, enhancers and 5’ and 3’ UTR. Neutral (negative) examples include more than 14 millions of SNVs in the non-coding regions of the reference human genome differing, according to high confidence alignment regions, from the ancestral primate genome sequence inferred on the basis of the Ensembl Enredo-Pechan-Ortheus whole-genome alignments of six primate species [39], and not including variants present in the most recent 1000 Genomes data [6] with frequency higher than 5% to remove variants that had not been exposed for sufficiently long time to natural selection. The imbalance between positive (mutations responsible for a Mendelian disease) and negative SNVs amounts to about 1:36,000. The 26 features associated to each SNV are genomic attributes ranging from G/C content, population-based features, to conservation scores, transcription and regulation annotations (for more details, see [29]).

We finally analyzed GWAS (Genome Wide Association Studies) data to detect 2115 regulatory GWAS hits downloaded from the National Human Genome Research Institute (NHGRI) GWAS catalog [40], and a set of negatives obtained by randomly sampling 1/10 of the negative examples of the Mendelian data set, according to the same experimental set-up described in [30], thus resulting in an imbalance between negative and positive examples of about 1 : 700. We predicted chromatin effect features directly from the DNA sequence using DeepSEA convolutional networks [18]: in this way we obtained 1842 features for each SNV, as described in [30], and we used that features to train *parSMURF*.

Table 2 summarizes the main characteristics of both the synthetic and genomic data used in our experiments.

**Table 2.** Summary of the datasets used in the experiments. Datasets are highly imbalanced towards the negative class.

| Name | N. of samples | N. of features | N. of positive samples | Imbalance ratio |
| --- | --- | --- | --- | --- |
| synth_1 | $10^6$ | 30 | 400 | 1 : 2500 |
| synth_2 | $10^7$ | 30 | 400 | 1 : 25000 |
| synth_3 | $5 \times 10^7$ | 30 | 1000 | 1 : 50000 |
| Mendelian | 14.755.605 | 26 | 406 | 1 : 36300 |
| GWAS | 1.477.630 | 1842 | 2115 | 1 : 700 |

### *parSMURF* scales nicely with synthetic & genomic data

Experiments reported in this section follow the classic experimental setup for the evaluation of the performances of parallel algorithms [41]. In particular, since our executions employ multiple computer processors (CPU) concurrently, we use speedup and efficiency to analyze the algorithm performances by measuring both the sequential and parallel execution times.

By denoting with *T*_*s*_ the run-time of the sequential algorithm and with *T*_*p*_ the run-time of the parallel algorithm executed on *P* processors, the speedup and efficiency are defined respectively as:

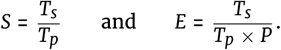

#### Speed-up and efficiency analysis with synthetic data

##### Experimental set-up

For every synthetic dataset, we run *parSMURF*^1^ and *parSMURF*^*n*^ several times by varying the number of threads (for both the multi-core and multi-node versions) and the number of MPI worker processes assigned to the task (for the multi-node version only). More precisely the number of threads *n.thr* varied in *n.thr* ∈ {1, 2, 4} for synth_1 and synth_2 datasets, while for synth_3 *n.thr* ∈ {1, 2, 4, 8, 16, 20}. Moreover we considered a number of MPI processes *n.proc* in the range *n.proc* ∈ {1, 2, 4, 8} for synth_1 and synth_2, and *n.proc* ∈ {1, 2, 4, 6, 8, 10} for synth_3.

For each run we collected the execution time and evaluated the speed-up and efficiency: the *T*_*s*_ sequential time of formulas (1) and (2) has been obtained by running *parSMURF*^1^ with 1 thread, hence obtaining a pure sequential run.

All experiments were executed using a 10-fold cross validation setting. The learning hyper-parameters employed in each experiment are the following:

- synth_1: *nParts* = 128, *fp* = 1, *ratio* = 1, *k* = 5, *nTrees* = 128, *mTry* = 5;
- synth_2: *nParts* = 64, *fp* = 1, *ratio* = 1, *k* = 5, *nTrees* = 32, *mTry* = 5;
- synth_3: *nParts* = 128, *fp* = 1, *ratio* = 5, *k* = 5, *nTrees* = 10, *mTry* = 5

For each synthetic data set we run experiments considering all the different numbers of threads *n.thr* for *parSMURF*^1^, while for *parSMURF*^*n*^ we run different hyper-ensembles considering all the possible combinations of *n.thr* and *n.proc*.

##### Results and discussion

Figure 5 reports the results for the batch of executions with the synth_1 and synth_2 datasets. Results are grouped by the number of MPI working processes (each line represents three runs obtained by keeping the number of MPI processes fixed and by varying the number of threads per process).

**Figure 5.**
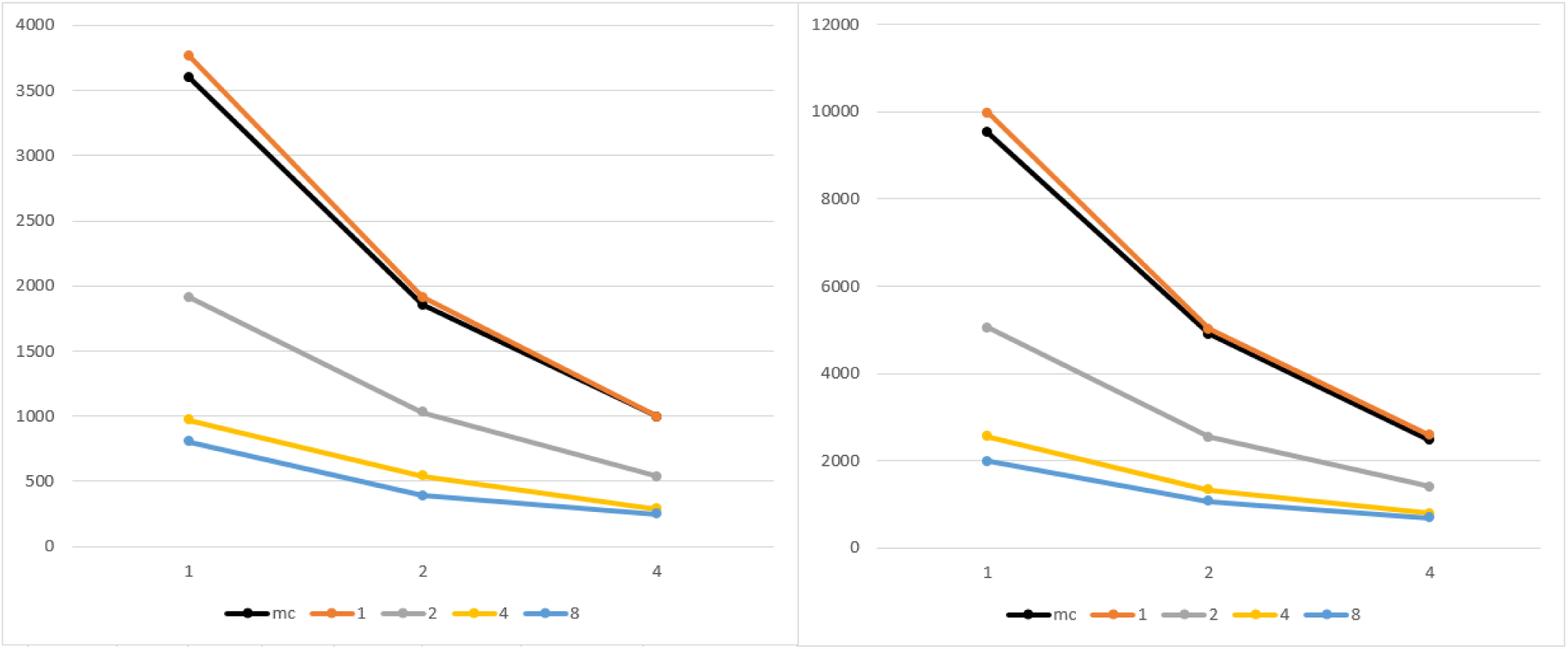
Execution time of *parSMURF*^1^ and *parSMURF*^*n*^ on the synthetic datasets synth_1 (left) and synth_2 (right). On x axis, the number of threads for each MPI process; on y axis, execution time in seconds; experiments are grouped by the number of MPI processes. Black line is the multi-thread version, while orange, gray, yellow and light blue are the MPI version with 1, 2, 4, and 8 MPI processes.

Both graphs show that the multi-core and the multi-node implementation of *parSMURF* introduce a substantial speed-up with respect to the sequential version (the point in the black line with 1 thread in the abscissa). Note that in Figure 5 the black line represents *parSMURF*^1^, while the orange line *parSMURF*^*n*^: their execution time is very similar since both use the same overall number of threads, with a small overhead for the MPI version due to the time needed to setup the MPI environment. Table 3 shows that the speed-up achieved by *parSMURF*^*n*^ is quasi-linear with respect to the overall number of “aggregated threads” (i.e. the product *n.thr* × *n.proc*) involved in the computation, at least up to 16 threads. By enlarging the number of “aggregated threads” we have a larger speed-up, but a lower efficiency, due to the lower number of parts of the partition assigned to each thread, and to the larger time consumed by the MPI data intercommunication.

**Table 3.** Execution time and speed-up of *parSMURF*^*n*^ with synth_1 and synth_2 datasets. Threads are counted as “aggregated” in the sense that they are the product of the number of MPI processes with the number of threads for each process. All executions have been performed with a 10-fold cross-validation setting.

| Aggregated threads | *synth_1* time (s) | *synth_1* speed-up | *synth_2* time (s) | *synth_2* speed-up |
| --- | --- | --- | --- | --- |
| 1 | 3768.59 | – | 9981.82 | – |
| 2 | 1910.19 | 1.97 | 5020.18 | 1.99 |
| 4 | 571.56 | 3.88 | 2539.10 | 3.93 |
| 8 | 542.35 | 6.95 | 1329.74 | 7.51 |
| 16 | 288.82 | 13.05 | 788.31 | 12.66 |
| 32 | 248.84 | 15.18 | 686.41 | 14.54 |

Results with the synthetic dataset synth_3, that includes 50 million examples, confirm that *parSMURF* scales nicely also when the size of the data is significantly enlarged. Indeed Figure 6 (left) shows that by increasing the number of processes and threads we can obtain a significant reduction of the execution time. These results are confirmed by grouping the execution time with respect to the “aggregated” number of threads, i.e. the product *n.thr* × *n.proc* (Figure 6 (right)).

**Figure 6.**
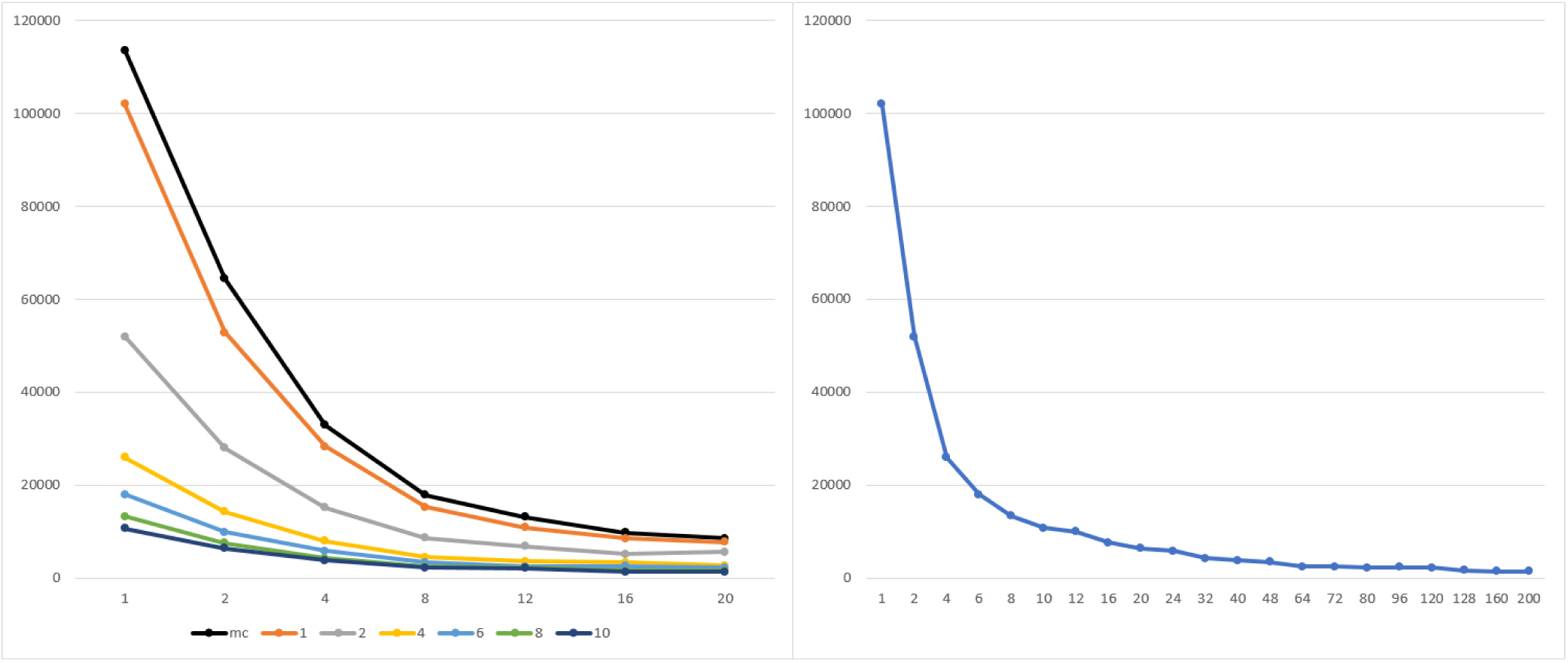
Execution time of *parSMURF*^1^ and *parSMURF*^*n*^ on the synthetic dataset synth_3. Left: on x axis, the number of threads for each MPI process; on y axis, execution time in seconds. Experiments are grouped by the number of MPI processes. Black line is the multi-thread version, while orange, gray, yellow, light blue, green and blue are the MPI version with 1, 2, 4, 6, 8 and 10 MPI processes. Right: results are grouped by total number of threads (*n.thr* × *n.proc*). When a combination is obtainable in more than one way only the best time is considered.

Figure 7 shows the speed-up (left) and efficiency (right) obtained with this dataset; results are again grouped by the “aggregated” number of threads. Note that with this large dataset we can obtain a larger speed-up, even if, as expected, at the expenses of the overall efficiency.

**Figure 7.**
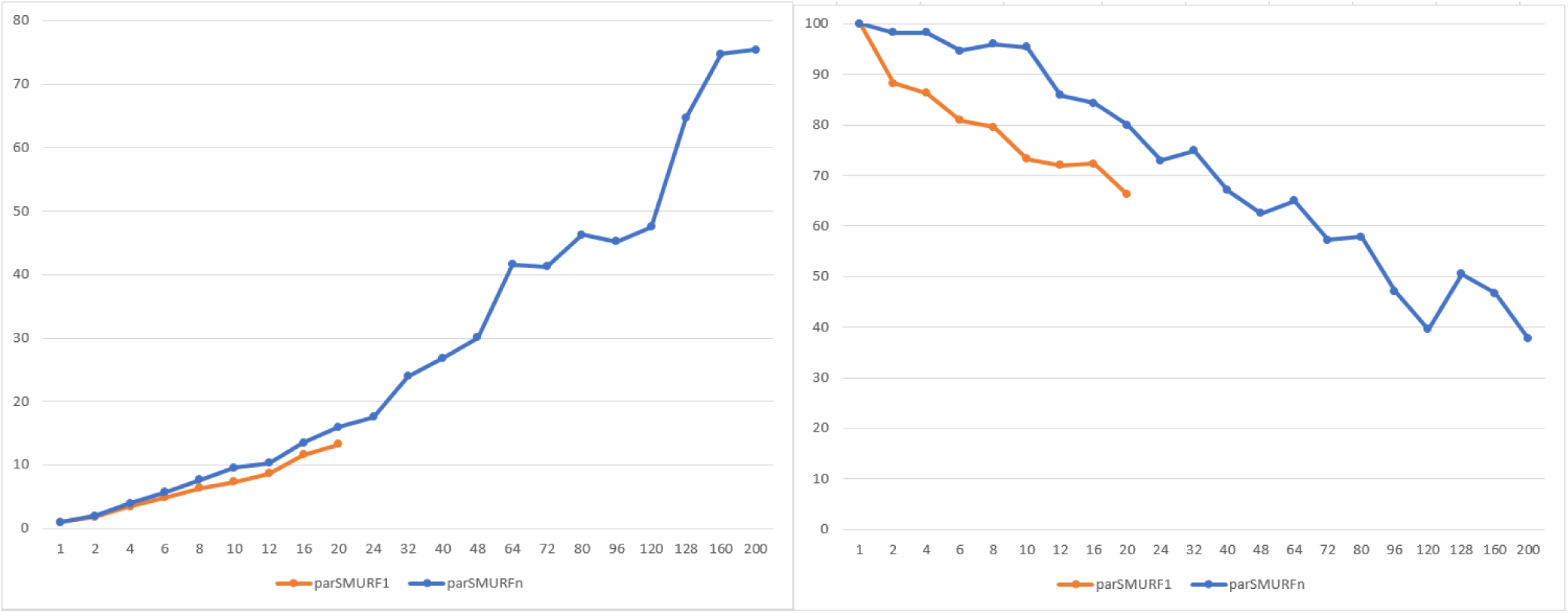
Left: Speed-up of *parSMURF*^1^ and *parSMURF*^*n*^ on the synthetic dataset synth_3. On x axis, the “aggregated” number of threads; on y axis, speed-up. Right: efficiency of *parSMURF* with the synthetic dataset synth_3. On x axis, the “aggregated” number of threads; on y axis, efficiency in percentage. Blue line refers to *parSMURF*^*n*^, orange line to *parSMURF*^1^.

Different research works showed contradictory results about the comparison of the performance of pure multiprocess MPI, pure multithread OpenMP or hybrid MPI-OpenMP implementations of the same algorithm, showing that several factors, such as algorithms, data structures, data size, hardware resources, MPI and OpenMP library implementations, influence their performances [42, 43, 44, 45, 46, 47, 48, 49].

Regarding our experiments, from Figure 6 and 8, we can notice how, in some cases, a pure MPI implementation may outperform an heterogeneous MPI-multithread or a pure OpenMP-multithread implementation. However, more in general, *parSMURF*^*n*^ allows a higher degree of parallelism, thus resulting in a larger speed-up and efficiency with respect to the pure multithread *parSMURF*^1^ (Figure 7 and 9).

**Figure 8.**
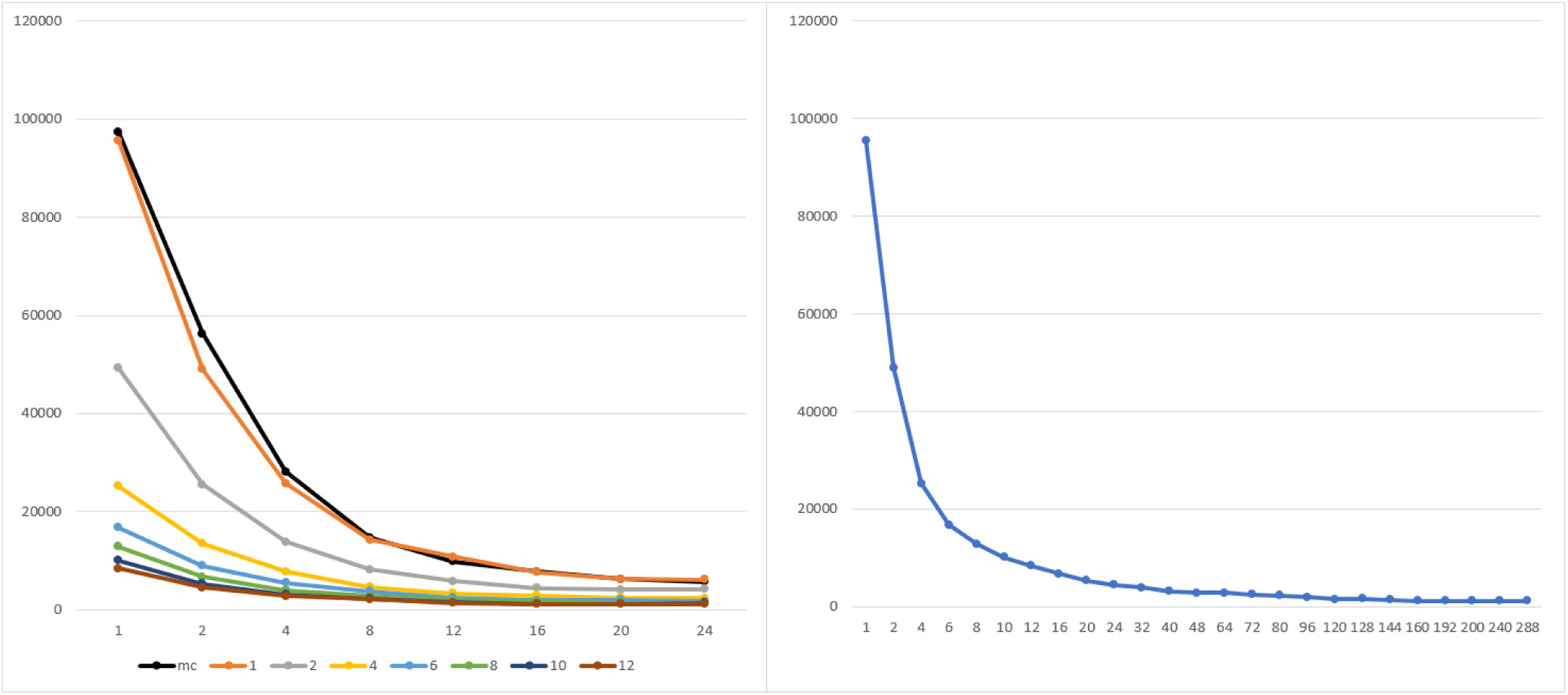
Execution time of *parSMURF*^1^ and *parSMURF*^*n*^ on the Mendelian data set. Left: on x axis, the number of threads for each MPI process; on y axis, execution time in seconds. Experiments are grouped by number of MPI processes. Black line is the multi-thread version, while orange, gray, yellow, light blue, green, blue and brown are the MPI version with 1, 2, 4, 6, 8, 10 and 12 MPI processes. Right: results are grouped by total number of threads (*n.thr* × *n.proc*).

**Figure 9.**
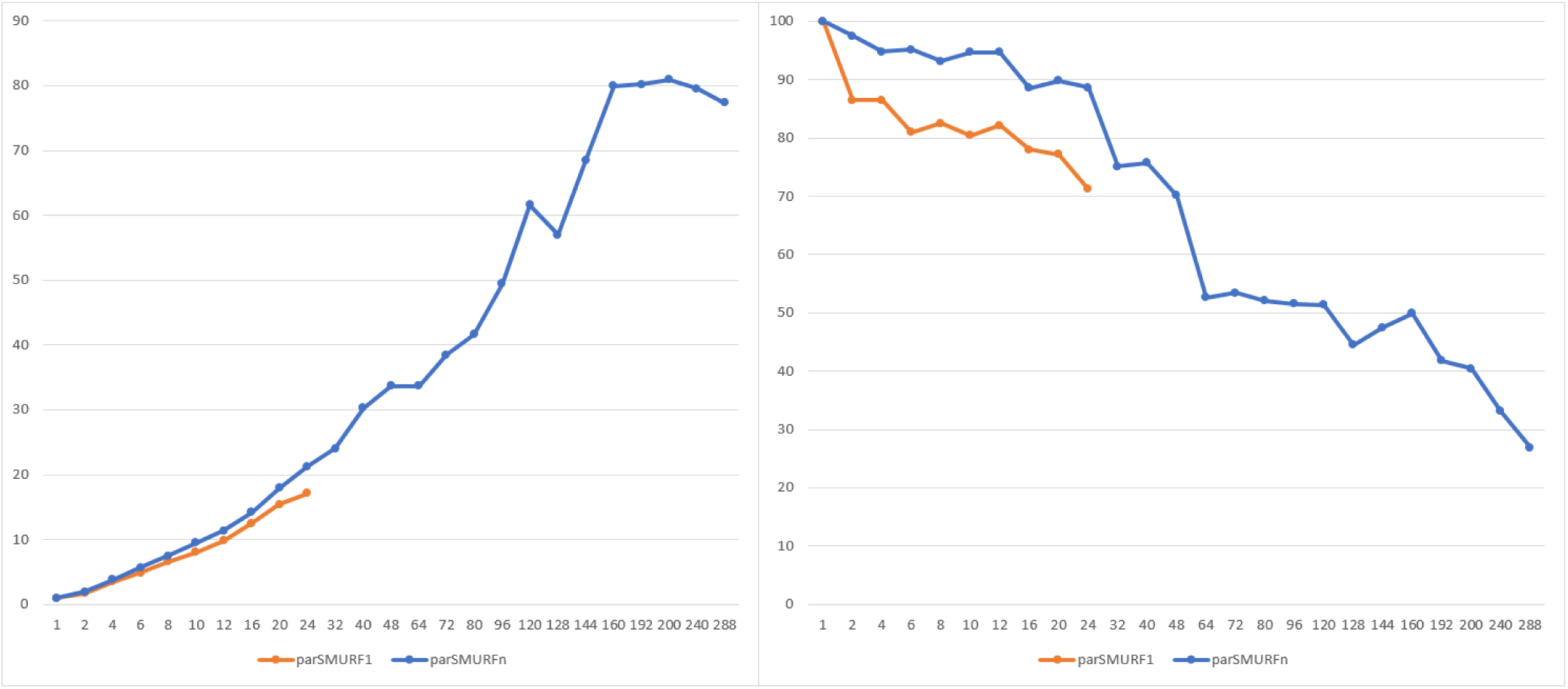
Left: Speed-up of of *parSMURF*^1^ and *parSMURF*^*n*^ with the Mendelian dataset. On x axis, the “aggregated” number of threads; on y axis, speed-up. Right: efficiency of *parSMURF* with the Mendelian dataset. On x axis, the “aggregated” number of threads; on y axis, efficiency in percentage. Blue line refers to *parSMURF*^*n*^, orange line to *parSMURF*^1^.

#### Speed-up and efficiency analysis with genomic data

To show how *parSMURF* performs in term of speed-up and efficiency on a real genomic dataset, we carried out the same batch of experiments as in the previous section using this time the Mendelian dataset.

Figure 8 shows the execution time of *parSMURF*^1^ and *parSMURF*^*n*^ as a function of the “aggregated number of threads”, i.e. the product of the number of MPI processes and the number of threads per process. As expected, results show a substantial decrement in execution time with respect to the number of the aggregated threads.

Figure 9 shows the speed-up and efficiency of *parSMURF*: on x axis of both graphs, threads are counted as “aggregated”, that is the total number of threads is computed by multiplying the number of processes by the number of threads assigned to each process. For the evaluation of speed-up and efficiency, *parSMURF*^1^ with only one computing thread has been used as reference for obtaining the computation time of the sequential version.

The maximum speed-up of *parSMURF*^1^ is about 17×, with the execution time decreasing from 97287 seconds of the sequential version to 5695 seconds of the multi-threaded version using 24 cores. The speed-up of *parSMURF*^*n*^ is even better, with a maximum speed-up of 80× (1181 seconds execution time) obtained using 10 MPI processes with 20 computing threads each. The graph shows that both *parSMURF*^1^ and *parSMURF*^*n*^ follow the same trend in the increment of the speed-up, but the multi-thread version is limited to 24 threads (each one assigned to a different core), while *parSMURF*^*n*^ continues this trend up to 288 threads, reaching a speed-up saturation level of 80×. As just observed with synthetic data (Figure 7), the efficiency tends to decrease with the number of aggregated threads.

Summarizing both experiments with synthetic and genomic data show that *parSMURF* scales nicely with large data and achieves a significant speed-up that allows its application to big data analysis and to the fine tuning of learning parameters.

### Auto-tuning of learning parameters improves prediction of pathogenic non-coding variants

The speed-up introduced by *parSMURF* allows the automatic fine tuning of its learning parameters to improve predictions on real genomic data. Indeed, as preliminarily shown in [33], fine tuning of *hyperSMURF* learning parameters can boost the performance on real data.

To this end we run *parSMURF*^*n*^ on the Cineca Marconi cluster using auto-tuning strategies to find the best learning parameters for both the prediction of pathogenic non-coding SNVs in Mendelian diseases and for the prediction of GWAS hits that overlap with a known regulatory element.

We compared the auto-tuned results only with those obtained with the default learning parameters of *hyperSMURF*, since our previous studies showed that *hyperSMURF* outperformed other methods, such as CADD [14], GWAVA [27], Eigen [19] and DeepSea [18] with both Mendelian diseases and GWAS hits data [30], and, above all, since we are more interested in showing a proof-of-concept of the fact that auto-tuning of learning parameters may lead to better performances in a real genomic problem.

#### Experimental set-up

Generalization performances have been evaluated through an external 10-fold “cytogenetic band-aware” cross-validation (CV) setting. This CV technique assures that variants occurring nearby in a chromosome (i.e. in the same cytogenetic band) do not occur jointly in the training and test sets and thereby biasing results, since nearby variants may share similar genomic features [30]. Learning parameters were selected through a grid search realized through a 9-fold internal CV, that is for each of the 10 training sets of the external cross-validation, their 9 ‘cytogenetic band-aware” folds have been used to select the best learning parameters and to avoid putting contiguous variants located in the same cytoband both in training and in the validation set.

This experimental set-up is computationally demanding, but by exploiting the different levels of parallelism available for *parSMURF*^*n*^ we can obtain a sufficient speed-up to experiment with different hyper-ensembles having different sets of learning parameters.

Performances of the prediction are evaluated via the Area Under the Receiver Operator Characteristic Curve (AUROC) and the Area Under the Precision-Recall Curve (AUPRC). Since data are highly unbalanced, we outline that it is well-known that in this context AUPRC provides more informative results [50, 51].

#### Improving predictions of pathogenic Mendelian variants

We at first executed *hyperSMURF* with default parameters (specifically: *nParts* = 100, *fp* = 2, *ratio* = 3, *k* = 5, *nTrees* = 10 and *mTry* = 5) in a context of 10-fold cytogentic-band aware CV, as this experiment is used as reference for the next steps.

We tested the auto-tuning feature by performing a grid search over the hyper-parameter space ℋ_*g*_ defined in Table 4, central column. Such hyperspace provides 576 possible hyper-parameter combinations *h* ∈ ℋ_*g*_. Then, we applied the auto-tuning strategy based on the Bayesian optimizer, by defining the hyper-parameter space ℋ_*b*_ as in Table 4, right column.

**Table 4.** Hyper-parameter spaces for grid search (ℋ_*g*_) and Bayesian optimizer (ℋ_*b*_) used for the auto-tuning on the Mendelian dataset.

| | $\mathcal{H}_g$ | $\mathcal{H}_b$ |
| --- | --- | --- |
| nParts | {10, 50, 100, 300} | [10, 300] |
| fp | {1, 2, 5, 10} | [1, 10] |
| ratio | {1, 2, 5, 10} | [1, 10] |
| k | {5} | {5} |
| nTrees | {10, 20, 100} | [10, 100] |
| mtry | {2, 5, 10} | [2, 10] |

To fully exploit the scalability of *parSMURF*, we launched the grid search with the following configuration: 10 instances of *parSMURF*^*n*^, one for each fold of the external CV, each one having 10 worker processes, with 6 dedicated threads for processing the different parts of the partition plus further 4 threads for each random forest training and testing. Hence, for the gridsearch, we used a total of 2400 CPU cores. Since the Bayesian auto-tuning procedure is less computationally intensive, we chose a more conservative approach on resource utilization for this experimental set-up: we launched one instance of *parSMURF*^*n*^ having 24 worker processes with 16 threads for the partitions and one for the random forest training and testing. Folds of the external CV are processed sequentially. Therefore, for the Bayesian optimization set-up we used 384 CPU cores.

At the end of this phase, for each optimization strategy, *parSMURF* returns the best hyper-parameter combination for each fold. We then executed 10 repetitions of the external CV using the default parameters, 10 using the best hyper-parameters found by the grid search and 10 using the best hyper-parameters found by the Bayesian optimizer. Performances in terms of AUROC and AUPRC were measured for each repetition and then averaged.

Performance improvements relative to the above parameter tuning experiments and their execution times are summarized in Table 7. Results in columns 4 and 5 show a significant improvement in the prediction performances in terms of AUPRC using both optimization strategies (Wilcoxon rank sum test, α = 10^−6^). On the other hand, AUROC is very high in all the experiments, confirming that this metric is not sufficient for the evaluation of prediction performances in the context of highly unbalanced datasets. Supplementary figures S1 and S2 show the computer ROC and precision-recall curves of both *hyperSMURF* and *parSMURF*. Also, the Bayesian optimizer proves to be effective in both improving the prediction performances and reducing the computational time: although slightly lower, predictions are comparable to the grid search, but they are obtained at a fraction of the computational power required by the latter. As a matter of fact, the CPU time required by the entire grid search counted more than 120*k* hours, compared to 16*k* hours used by the Bayesian optimization strategy.

**Table 7.**
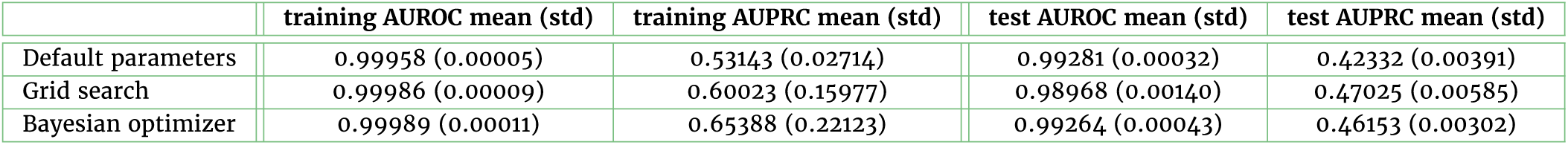
Summary of performance improvements obtained by *parSMURF* by tuning the learning parameters on the Mendelian dataset. Results are averaged across 10 repetitions of the 10-fold cytoband-aware cross-validation. “AUROC avrg” and “AUPRC avrg” are averaged across the 10 folds; standard deviation in brackets. Columns 2 and 3 report AUROC and AUPRC metrics on the training set, columns 4 and 5 report the same metrics evaluated on the test set. Default parameters: nParts 100, fp 2, ratio 3, nTrees 10, mTry 5.

|  | training AUROC mean (std) | training AUPRC mean (std) | test AUROC mean (std) | test AUPRC mean (std) |
| --- | --- | --- | --- | --- |
| Default parameters | 0.99958 (0.00005) | 0.53143 (0.02714) | 0.99281 (0.00032) | 0.42332 (0.00391) |
| Grid search | 0.99986 (0.00009) | 0.60023 (0.15977) | 0.98968 (0.00140) | 0.47025 (0.00585) |
| Bayesian optimizer | 0.99989 (0.00011) | 0.65388 (0.22123) | 0.99264 (0.00043) | 0.46153 (0.00302) |

Table 7 reports average AUROC and AUPRC measured on the training sets: results show that the ratio between training and test AUROC or AUPRC is quite similar between *hyperSMURF* and *parSMURF*, and even if, as expected, results on the training sets are better, they are comparable with those obtained on the test data. These results show that performance improvement is not due to overfitting, but to a proper choice of the hyper-parameters well-fitted to the characteristics of the problem.

To assess whether the difference in performance between *hyperSMURF* and *parSMURF* can be related to their different capacity of selecting the most informative features, we measured Spearman correlation between both *hyperSMURF* and *parSMURF* scores with each of the 26 features used to train the hyper-ensembles for all the examples of the dataset. Table S3 in Supplementary Information reports the correlation between the true labels of the examples and the predictions obtained by *hyperSMURF* using the default set of hyper-parameters, *parSMURF* with the default, grid optimized and Bayesian optimized set of hyper-parameters. We found that *hyperSMURF* and *parSMURF* achieved very similar Spearman correlation on each feature (the Pearson correlation between the vectors of Spearman correlations of *hyperSMURF* and *parSMURF* is about 0.98). Both *hyperSMURF* and *parSMURF* obtained the largest Spearman correlation coefficients for features related to the evolutionary conservation of the site (e.g. vertebrate, mammalian and primate PhyloP scores) and for some epigenomic features (histone acetylation, methylation and DNAse hypersensitivity). Again, these results show that it is unlikely that the improved performance of *parSMURF* can be explained through its better capacity of selecting the most informative features, but instead by its capacity of auto-tuning its learning hyper-parameters and its capacity to find a model that better fits the data.

In addition, in Table 6 some examples of pathogenic variants which have been ranked remarkably better by *parSMURF* than *hyperSMURF* are reported. Further details about these variants are shown in Table S6 of Supplementary Information.

**Table 6.** Examples of pathogenic Mendelian variants better ranked by *parSMURF* with respect to *hyperSMURF*. The first two columns report the chromosomal coordinates, while the last two the difference in ranking between respectively *parSMURF* with grid search (ℋ_*g*_) and with Bayesian optimizer (ℋ_*b*_) with respect to *hyperSMURF*. The larger the absolute difference, the higher the improvement (see also Table S6 in Supplementary for more information).

| chr | pos | hS rank - $\mathcal{H}_g$ rank | hS rank - $\mathcal{H}_b$ rank |
| --- | --- | --- | --- |
| 1 | 100661453 | 2308597 | 169786 |
| 3 | 12421189 | 663054 | 421027 |
| X | 138612889 | 194290 | 111499 |
| 13 | 100638902 | 70175 | 69069 |
| 6 | 118869423 | 63078 | 55789 |
| 16 | 31202818 | 50539 | 103623 |
| 12 | 121416444 | 21848 | 65773 |

#### *Prediction performances of* parSMURF *with an independent Mendelian test set*

We collected novel Mendelian pathogenic variants by adding 64 newly positive (pathogenic) non-coding variants manually annotated according to a comprehensive literature review. We included only those variations and publications judged to provide plausible evidence of pathogenicity (Supplementary Table 7). As negatives we used common variants downloaded from NCBI [52], i.e. variants of germline origin and having a minor allele frequency (MAF) ≥ 0.01 in at least one major population, with at least two unrelated individuals having the minor allele, where major populations are those included in the 1K genome project [53]. We selected only those variants that lie in non-coding regions using Jannovar [54]. The final number of negatives (about 3 millions of examples) has been randomly sampled in such a way that the ratio positives/negative in the original and in the new Mendelian data set used for validation is approximately the same. Both the positive and negative examples have been annotated with the same 26 genomic and epi-genomic features used for the original Mendelian data set. We trained *hyperSMURF* and *parSMURF* on the overall original Mendelian data set and then we tested the resulting models on the unseen separated new Mendelian data set used for validation. Since the new positive set also contains small insertions and deletions, similarly to [29], to predict the pathogenicity of the deletions, we used the maximum score of any nucleotide included in the deleted sequence, while for insertions we used the maximum score computed for the two nucleotides that surround the insertion. Results with the independent Mendelian test set show that the models are able to obtain relatively high AUPRC results, especially when *parSMURF* is applied, showing that our models can nicely generalize. Also with this new independent data set *parSMURF* significantly outperforms *hyperSMURF* (Table 5).

**Table 5.** *hyperSMURF* and *parSMURF* (with Grid Search and Bayesian optimization) prediction performances obtained over a fully independent Mendelian test set composed by newly annotated pathogenic variants (positive examples) and common neutral variants (negative examples).

| Model | AUROC | AUPRC |
| --- | --- | --- |
| *hyperSMURF* | 0.945216 | 0.098153 |
| *parSMURF*- Grid Search | 0.941026 | 0.409067 |
| *parSMURF*- Bayesian optimization | 0.928158 | 0.192568 |

#### Improving predictions of GWAS hits

A similar experimental setup has been employed for improving the predictions of GWAS hits. At first we executed *parSMURF* with the default parameters as reference for the next batches of experiments. Then, we tested the auto-tuning feature by performing a grid search over the hyper-parameter space ℋ_*g*_ defined in Table 8, central column. Such hyperspace provides 256 possible hyper-parameter combinations *h* ∈ ℋ_*g*_. Next, we tested the Bayesian optimizer by defining the hyper-parameter space ℋ_*b*_ as in Table 8, right column.

**Table 8.** Hyper-parameter space for Grid search and Bayesian Optimization used for auto-tuning *parSMURF* on the GWAS data set.

|  | Grid-search | Bayesian-opt. |
| --- | --- | --- |
| *nParts* | {10, 20, 30, 40} | [10, 40] |
| *fp* | {1, 2, 5, 10} | [1, 10] |
| *ratio* | {1, 2, 5, 10} | [1, 10] |
| *k* | {5} | {5} |
| *nTrees* | {10, 20, 50, 100} | [10, 100] |
| *mtry* | {30} | {30} |

Results are shown in Table 9. As for the Mendelian dataset, AUROC is very high in all experiments. On the other hand, test results show a significant increase of AUPRC with both auto-tuning strategy, with the grid search leading a better outcome than the Bayesian optimizer. Supplementary figures S3 and S4 show the ROC and precision-recall curves of *hyperSMURF* and *parSMURF*.

**Table 9.**
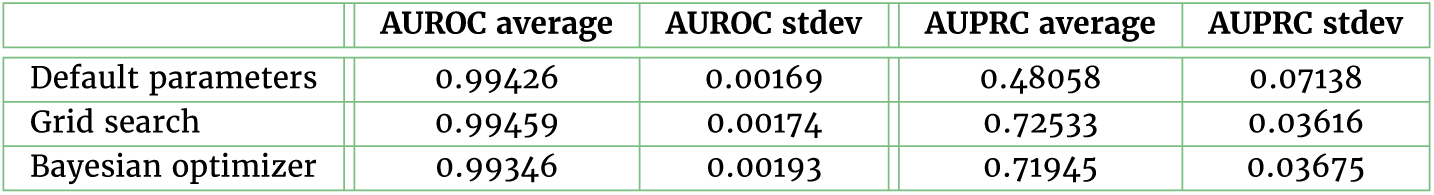
Summary of the performance improvements obtained by *parSMURF* by tuning its learning parameters with the GWAS Catalog dataset. Results are averaged across 10 repetitions of the 10-fold cytoband-aware cross-validation. “AUROC avrg” and “AUPRC avrg” are averaged across the 10 folds. Default par: nParts 100, fp 2, ratio 3, nTrees 10, mTry 5.

|  | AUROC average | AUROC stdev | AUPRC average | AUPRC stdev |
| --- | --- | --- | --- | --- |
| Default parameters | 0.99426 | 0.00169 | 0.48058 | 0.07138 |
| Grid search | 0.99459 | 0.00174 | 0.72533 | 0.03616 |
| Bayesian optimizer | 0.99346 | 0.00193 | 0.71945 | 0.03675 |

These results further show that fine tuning of learning parameters is fundamental to significantly improve prediction performances, showing also that *parSMURF* is a useful tool to automatically find “near-optimal” learning parameters for the prediction task under study.

#### Assessment of the effect on prediction performance of the variants imbalance across regulatory regions

As recently pointed out in [28], pathogenic scores predicted by several state-of-the-art methods are biased towards some specific regulatory region types. Indeed also with Mendelian and GWAS data the positive set of variants is located in different functional non-coding regions (like 5’UTR, 3’UTR or Promoter) and is not evenly distributed over them. This is also the case for the negative set (see Supplementary Table S4 and S5). Because of this imbalance, performances of different categories are different as already mentioned by Smedley et al. for the ReMM score on the Mendelian data [29]. It is possible that our *parSMURF* parameter optimization will favor different categories because of the number of available positives and the different imbalance between positives and negatives across different genomic regions. To show that the optimization is robust to this characteristic of the data we compared performances on each genomic category before and after parameter optimization. Variant categories have been defined through Jannovar [54] using the RefSeq database.

Then we retrained and re-optimized the parameters on a training set using cytoband-aware cross-validation, where all categories have the same imbalance by subsampling negatives to the smallest imbalance of the categories. More in detail, we used the following strategy: (1) sub-sample the negatives to the same imbalance in all categories. Mark the variant if it is in this new subset; (2) partition the whole dataset into 10 folds as done previously; (3) for each training step select only the previously marked variants of the 9 training folds; (4) sub-sample the test set using the same categorization and ratios as in (1). To take into account the variability between runs, we repeated this process 10 times for the Mendelian dataset and 5 times for the GWAS dataset. Using this strategy both training and test sets are equally “per region balanced”, so that category unbalance is kept under control and we can correctly evaluate whether our approach may unnaturally inflate predictions towards a specific region due to the original per-region imbalance of both datasets.

Results are shown in Supplementary Figures S5 and S6: for all variant categories we see a performance gain or similar performance in *parSMURF* with respect to *hyperSMURF* for both the Mendelian and GWAS data set, suggesting that *parSMURF* is robust to the categorical composition of the variants. Moreover in the “per region balanced” setting AUPRC results are systematically better with the Mendelian data set (Supplementary Figures S5) and quite always better or comparable with the GWAS data (Figure S6). These experimental results show that both *hyperSMURF* and *parSMURF* can properly handle different imbalances of variant categories, by using “smart” balancing techniques on the training set able to both balancing and at the same time maintaining a large coverage of the available training data. The increase of performance of *parSMURF* with respect to *hyperSMURF* is not driven by the under- or over-representation of variants belonging to a particular region type, but by its capacity of automatically fine-tuning the set of its hyper-parameters, according to the given task at hand.

#### Analysis of the hyper-parameters

Since we adopted cross-validation techniques to estimate the generalization performance of the models, we averaged the best parameters values separately estimated for each fold, in order to obtain a single set of optimal parameters. Tables S1 and S2 of the Supplementary Information show the sets of best hyper-parameters found by both the optimization techniques with the Mendelian and GWAS datasets.

Of the 6 hyper-parameters, we noticed that *nParts, fp* and *ratio* are the main factors that drive the performance improvement. *Fp* and *ratio* hyper-parameters provide the rebalancing of the classes. A larger *fp* value translates into a larger number of positive examples generated through the SMOTE algorithm (see Section Methods), thus reducing the imbalance between positive and negative examples in the training set: Tables S1 and S2 of the Supplementary show that enlarging the ratio of novel positive examples *parSMURF* improves results over *hyperSMURF*, and confirm that fine-tuned balancing techniques can improve the results. The *ratio* hyper-parameter controls the ratio between negative and positive examples of the training set. Results in Tables S1 and S2 show that values larger than the default ones improve performance, since in this way we can both reduce the imbalance between negatives and positives (for the Mendelian data sets we move from 36000 : 1 to 10 : 1, and for GWAS from 700 : 1 to 10 : 1), and at the same time we maintain a relatively large coverage of the negative data (in each partition negative examples are sampled in such a way to obtain ten negatives for each positive of the training set).

The results also show that a larger coverage of negative examples is obtained by incrementing the *nParts* hyper-parameter, since by increasing the number of partitions, less negatives are discarded. Moreover more random forests are trained thus improving the generalization capabilities of the hyper-ensemble. Finally, for the GWAS dataset, the *mtry* hyper-parameter plays a fundamental role in the increment of the performance, due to the high number of features of the dataset. Overall, the analysis of the hyper-parameters confirms that their fine tuning is fundamental to improve the performances of the hyper-ensemble.

## Conclusion

In this paper we presented *parSMURF*, a High Performance Computing tool for the prediction of pathogenic variants, designed to deal with the issues related to the inference of accurate predictions with highly unbalanced datasets. We showed that *hyperSMURF*, despite its encouraging results with different genomic data sets, suffers from two major drawbacks: a very demanding computing time and the need of a proper fine tuning of the learning parameters. The proposed *parSMURF* method provides a solution for both problems, through two efficient parallel implementations - *parSMURF*^1^ and *parSMURF*^*n*^ - that scale well with respectively multi-core machines and multi-node HPC cluster environments.

Results with synthetic datasets show that *parSMURF* scales nicely with large data sets, introducing a sensible speed-up with respect to the pure sequential version. Especially for large data sets, as expected, we should prefer the hybrid MPI-multi-thread version *parSMURF*^*n*^, while for relatively smaller datasets we can obtain a reasonable speed-up also with the pure multi-thread version *parSMURF*^1^ that can run also with a off-the-shelf laptop or desktop computer, by exploiting the multi-core architecture of modern computers.

*parSMURF* features two different and both effective strategies for the auto-tuning of the learning parameters: the first is based on an exhaustive grid search which proves to be effective in finding the best combination of hyper-parameters in terms of maximizing the AUPRC rating, but turns out to be very computing intensive. The other strategy is Bayesian optimization based and aims to find a near-optimal hyper-parameter combination in a fraction of time compared to the grid search strategy. Experimental results with Mendelian diseases and GWAS hits in non-coding regulatory regions show that *parSMURF* can enhance *hyperSMURF* performance, confirming that fine-tuning of learning hyper-parameters may lead to significant improvements of the results.

The high level of parallelism of *parSMURF*, its autotuning hyper-parameters capabilities and its easy-to-use software interface allow the user to apply this tool to ranking and classification problems characterized by highly imbalanced big data. This situation commonly rises up in Genomic Medicine problems, since only a small set of “positive” examples is usually available to train the learning machines. For this reason *parSMURF* can be a useful tool not only for the prediction of pathogenic variants, but also for any imbalanced ranking and classification problem in Genomic Medicine, provided that suitable big data are available for the problem at hand.

## Availability of source code and requirements

Project name: ParSMURF

Project home page: https://github.com/AnacletoLAB/parSMURF

GitHub repository: https://github.com/AnacletoLAB/parSMURF

SciCrunch RRID: `SCR_017560`

Operating system(s): Linux

Programming language: C++, Python 2.7

Requirements for *parSMURF*^1^: Multi-core x86-64 processor,

512 MB RAM, C++ compiler supporting OpenMP standard. Requirements for *parSMURF*^*n*^: Multi-core x86-64 processor, 1024 MB RAM, implementation of MPI library (i.e. OpenMPI or IntelMPI) installed on each node of the cluster, a reasonably fast interconnecting infrastructure.

License: GNU General Public License v3

## Availability of supporting data and materials

Datasets used for the assessment of scalability and prediction quality are available at the following page of the Open Science Foundation project: [55]. Supplementary Information with detailed experimental results are downloadable from the GigaScience website.

## Declarations

### List of abbreviations

AUPRC: Area Under the Precision-Recall Curve
AUROC: Area Under the Receiver-Operating-Characteristic Curve
CADD: Combined Annotation-Dependant Depletion
CV: Cross-Validation
FATHMM-MKL: Functional Analysis through Hidden Markov Models and Multiple Kernel Learning
G/C content: Guanine-Cytosine content
gkm-SVM: Gapped k-mer Support Vector Machine
GWAS: Genome-Wide Association Study
GWAVA: Genome Wide Annotation of Variants
ML: Machine Learning
MPI: Message Passing Interface
NGS: Next Generation Sequencing
OpenMP: Open Multi-Processing
RF: Random Forest
SLURM: Simple Linux Utility for Resource Management
SMOTE: Synthetic Minority Over-Sample technique
SNV: Single Nucleotide Variants
UTR: Untraslated Region

## Author’s Contributions

All the authors contributed to this paper. Conceptualization and methodology: A.P., G.V. Formal analysis: A.P., G.V., G.G,M.F. Data Curation and Investigation: M.S., M.R., D.D. Software: A.P., G.G. M.F. and L.C. Supervision: G.V., P.R. Validation: A.P., T.C. Funding acquisition: G.G., M.M., G.V. Writing - Original Draft Preparation: G.V., A.P., M.M. Writing - Review & Editing: all the authors.

