## Supplementary figures and tables for "*parSMURF*, a High Performance Computing tool for the genome-wide detection of pathogenic variants"

*GigaScience*, 2019, 1–9

doi: xx.xxxx/xxxx
Manuscript in Preparation
Supplementary material

SUPPLEMENTARY MATERIAL

### *parSMURF*, a High Performance Computing tool for the genome-wide detection of pathogenic variants – Supplementary Information

Alessandro Petrini[1], Marco Mesiti[1], Max Schubach[2,3], Marco Frasca[1], Daniel Danis[4], Matteo Re[1], Giuliano Grossi[1], Luca Cappelletti[1], Tiziana Castrignanò[5], Peter N. Robinson[4] and Giorgio Valentini[1]

[1]AnacletoLab – Dipartimento di Informatica, Università degli Studi di Milano, Italy and [2]Berlin Institute of Health (BIH), Berlin, Germany and [3]Charité – Universitätsmedizin Berlin, Berlin, Germany and [4]The Jackson Laboratory for Genomic Medicine, Farmington CT, USA and [5]CINECA, SCAI SuperComputing Applications and Innovation Department, Roma, Italy

5 Prediction performances (AUROC and AUPRC) for the Mendelian dataset, with both the Original imbalanced Mendelian data set and with the separated "per-region balanced" Mendelian data used for validation using *hyperSMURF* (hS) and *parSMURF* (pS) with Bayesian optimization hyper-ensembles (set of hyper-parameters found by the Bayesian optimizer: `nParts`: 36 - `fp`: 3 - `ratio`: 5 - `k`: 5 - `numTrees`: 55 - `mtry`: 5). Each group shows prediction performances divided by regulatory region type. "Global" group shows prediction performances of the entire dataset, disregarding the regulatory region type. AUROC and AUPRC have been averaged across "cytoband-aware" 10-fold cross validation. Results have been averaged across 10 repetitions of the same experimental setup (error bars represent standard deviation). "Original imbalanced" and "Per-region balanced" stands respectively for the original imbalanced setting and the balanced setting where the same ratio of positives versus negatives in both training and test sets is maintained in each regulatory region. . . . . . . . . . . . . . . . . . . . . . . . . 4
6 Prediction performances (AUROC and AUPRC) for the GWAS dataset, with both the Original imbalanced GWAS data set and with the separated "per-region balanced" GWAS data used for validation using *hyperSMURF* (hS) and *parSMURF* (pS) with Bayesian optimization hyper-ensembles (set of hyper-parameters found by the Bayesian optimizer: `nParts`: 10 - `fp`: 2 - `ratio`: 5 - `k`: 5 - `numTrees`: 50 - `mtry`: 30). Each group shows prediction performances divided by regulatory region type. "Global" group shows prediction performances of the entire dataset, disregarding the regulatory region type. AUROC and AUPRC have been averaged across "cytoband-aware" 10-fold cross validation. Results have been averaged across 5 repetitions of the same experimental setup (error bars represent standard deviation). "Original imbalanced" and "Per-region balanced" stands respectively for the original imbalanced setting and the balanced setting where the same ratio of positives versus negatives in both training and test sets is maintained in each regulatory region. . . . 5

#### List of Tables

1 Optimal sets of hyper-parameters returned by the optimizers embedded in *parSMURF* while training the model with the Mendelian dataset. Refer to Table 1 in the main paper for a brief explanation of each parameter. Default values are those used with *hyperSMURF*. As Grid and Bayesian optimizer returns the set of best hyper-parameters for each fold of the cross-validation, optimal values have been computed by averaging each parameter over the folds. . . . . . . . . . . . . . . . . . . . . . . . 6
2 Optimal sets of hyper-parameters returned by the optimizers embedded in *parSMURF* while training the model with the GWAS dataset. Refer to Table 1 in the main paper for a brief explanation of each parameter. Default values are those used with *hyperSMURF*. As Grid and Bayesian optimizer returns the set of best hyper-parameters for each fold of the cross-validation, optimal values have been computed by averaging each parameter over the folds. . . . . . . . . . . . . . . . . . . . . . . . 6
3 Spearman correlation between HyperSMURF and parSMURF scores for each of the 26 features of the Mendelian dataset. . . . . . . . . . . . . . . . . . . 6
4 Imbalance of the number of negative and positive examples across different regulatory region types in the Mendelian dataset. In column 4, the ratio between negative and positive samples; in bold, the lowest ratio used to under-sample the negatives for each regulatory region. The last column reports the number of down-sampled negatives for each regulatory region after applying the best ratio found in column 4. . . . . . . . . . . . . . 6
5 Imbalance of the number of negative and positive samples across different regulatory region types in the GWAS dataset. In column 4, the ratio between negative and positive samples; in bold, the lowest ratio used to under-sample the negatives for each regulatory region. The last column reports the number of down-sampled negatives for each regulatory region after applying the best ratio found in column 4. . . . . . . . . . . . . . . . . 7
6 Examples of pathogenic Mendelian single nucleotide variants where *parSMURF* sensibly outperformed *hyperSMURF*. The first two columns refer to the chromosomal coordinates of the variant. Ref and Alt to the reference and alternative allele; OMIM to the OMIM code of the associated Mendelian disease; Gene to the symbolic name of the target gene; PMID to the PubMed ID of the related publication; Region to the type of the regulatory region. The two last columns refer to the difference of ranking between respectively *parSMURF* with grid optimization and *parSMURF* with Bayesian optimization with respect to *hyperSMURF*. . . . . . . . . . . . . . . . . . . . . . . . 7
7 List of newly annotated pathogenic variants used as independent test set to assess the generalization capabilities of *parSMURF*. The first two columns refer to chromosomal coordinates, REF and ALT to the reference and alternative allele (including both SNVs, micro-insertions and micro-deletions), Region the type of regulatory region, Gene the symbolic name of the gene and PMID the PubMed ID of the related publication. . . . . . . 8

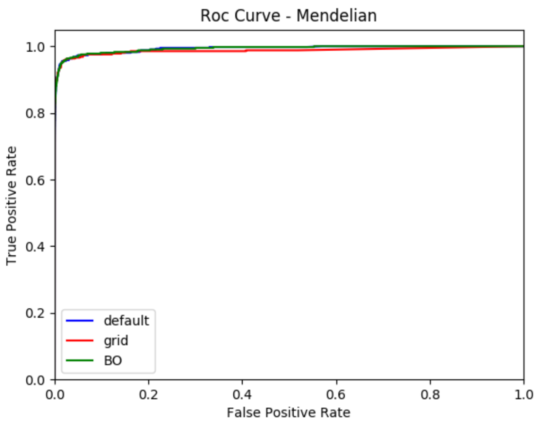

**Figure 1.** Plot of Receiver Operating Characteristic curve of the predictions for the Mendelian dataset using 3 sets of hyper-parameters: default (blue), best from the grid search (red) and best from the Bayesian optimizer (green)

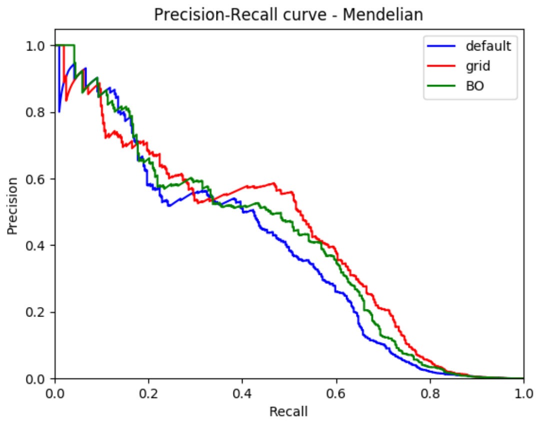

**Figure 2.** Plot of Precision-Recall curve of the predictions for the Mendelian dataset using 3 sets of hyper-parameters: default (blue), best from the grid search (red) and best from the Bayesian optimizer (green)

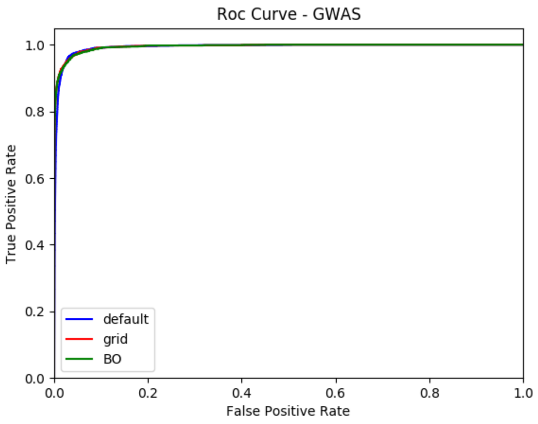

**Figure 3.** Plot of Receiver Operating Characteristic curve of the predictions for the GWASn dataset using 3 sets of hyper-parameters: default (blue), best from the grid search (red) and best from the Bayesian optimizer (green)

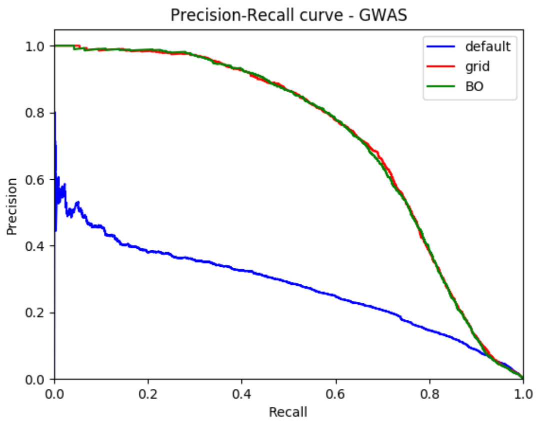

**Figure 4.** Plot of Precision-Recall curve of the predictions for the GWAS dataset using 3 sets of hyper-parameters: default (blue), best from the grid search (red) and best from the Bayesian optimizer (green)

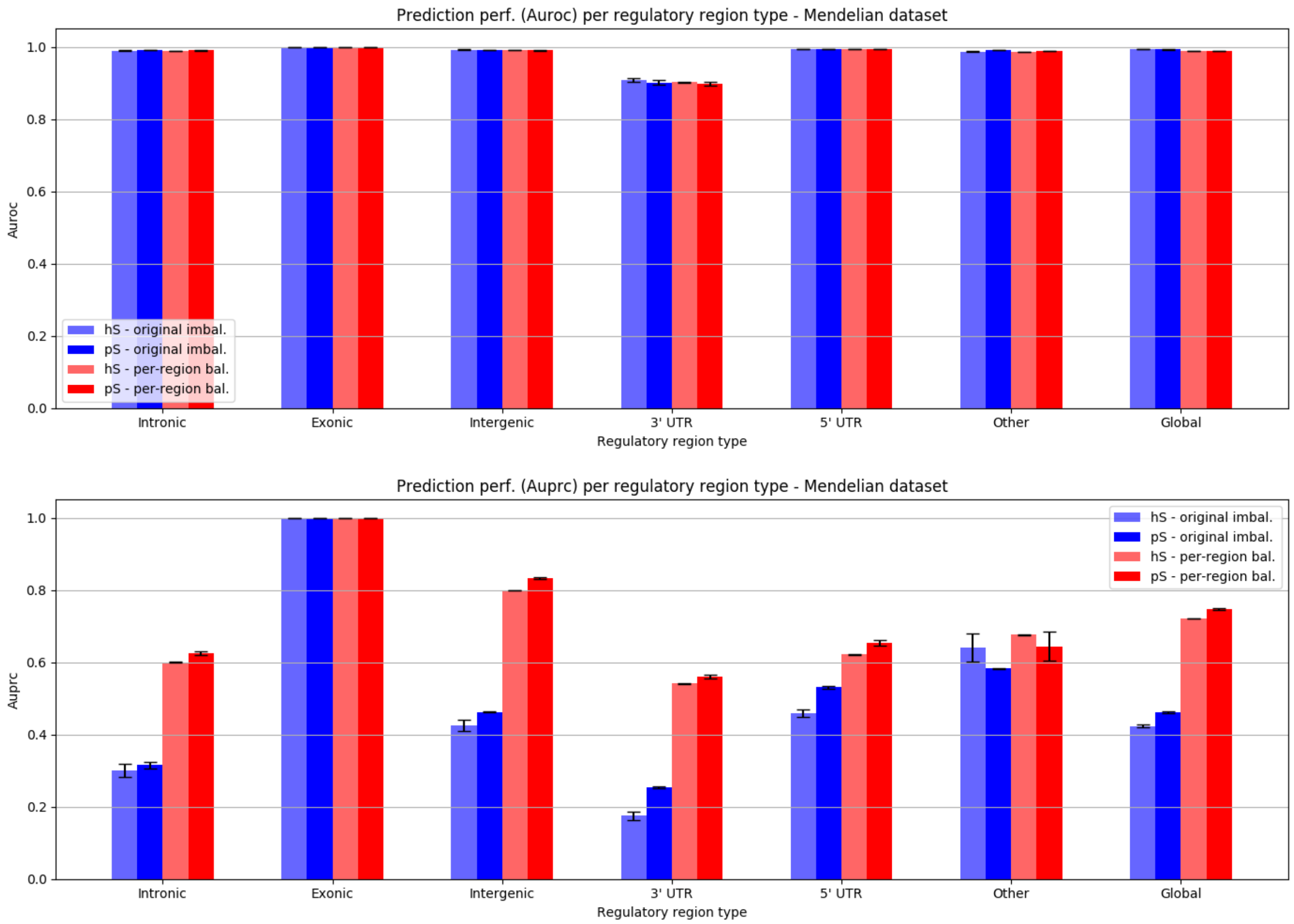

**Figure 5.** Prediction performances (AUROC and AUPRC) for the Mendelian dataset, with both the Original imbalanced Mendelian data set and with the separated "per-region balanced" Mendelian data used for validation using *hyperSMURF* (hS) and *parSMURF* (pS) with Bayesian optimization hyper-ensembles (set of hyper-parameters found by the Bayesian optimizer: `nParts`: 36 - `fp`: 3 - `ratio`: 5 - `k`: 5 - `numTrees`: 55 - `mtry`: 5). Each group shows prediction performances divided by regulatory region type. "Global" group shows prediction performances of the entire dataset, disregarding the regulatory region type. AUROC and AUPRC have been averaged across "cytoband-aware" 10-fold cross validation. Results have been averaged across 10 repetitions of the same experimental setup (error bars represent standard deviation). "Original imbalanced" and "Per-region balanced" stands respectively for the original imbalanced setting and the balanced setting where the same ratio of positives versus negatives in both training and test sets is maintained in each regulatory region.

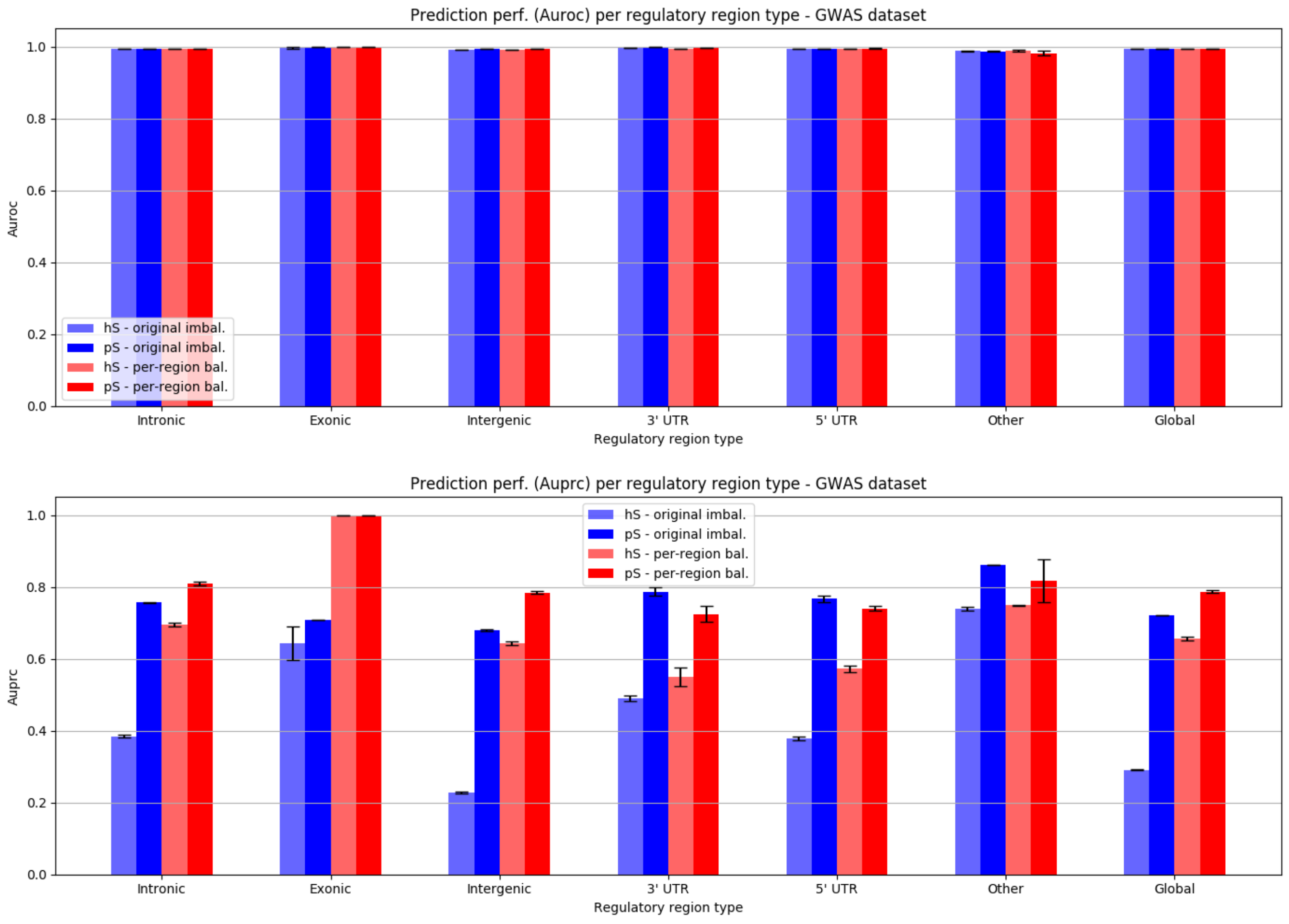

**Figure 6.** Prediction performances (AUROC and AUPRC) for the GWAS dataset, with both the Original imbalanced GWAS data set and with the separated "per-region balanced" GWAS data used for validation using *hyperSMURF* (hS) and *parSMURF* (pS) with Bayesian optimization hyper-ensembles (set of hyper-parameters found by the Bayesian optimizer: `nParts`: 10 - `fp`: 2 - `ratio`: 5 - `k`: 5 - `numTrees`: 50 - `mtry`: 30). Each group shows prediction performances divided by regulatory region type. "Global" group shows prediction performances of the entire dataset, disregarding the regulatory region type. AUROC and AUPRC have been averaged across "cytoband-aware" 10-fold cross validation. Results have been averaged across 5 repetitions of the same experimental setup (error bars represent standard deviation). "Original imbalanced" and "Per-region balanced" stands respectively for the original imbalanced setting and the balanced setting where the same ratio of positives versus negatives in both training and test sets is maintained in each regulatory region.

**Table 1.** Optimal sets of hyper-parameters returned by the optimizers embedded in *parSMURF* while training the model with the Mendelian dataset. Refer to Table 1 in the main paper for a brief explanation of each parameter. Default values are those used with *hyperSMURF*. As Grid and Bayesian optimizer returns the set of best hyper-parameters for each fold of the cross-validation, optimal values have been computed by averaging each parameter over the folds.

| | nParts | fp | ratio | k | nTrees | mtry |
|---|---|---|---|---|---|---|
| Default | 100 | 2 | 3 | 5 | 10 | 5 |
| Grid search | 260 | 7 | 10 | 5 | 50 | 4 |
| Bayesian optimizer | 74 | 8 | 10 | 5 | 50 | 5 |

**Table 2.** Optimal sets of hyper-parameters returned by the optimizers embedded in *parSMURF* while training the model with the GWAS dataset. Refer to Table 1 in the main paper for a brief explanation of each parameter. Default values are those used with *hyperSMURF*. As Grid and Bayesian optimizer returns the set of best hyper-parameters for each fold of the cross-validation, optimal values have been computed by averaging each parameter over the folds.

| | nParts | fp | ratio | k | nTrees | mtry |
|---|---|---|---|---|---|---|
| Default | 100 | 2 | 3 | 5 | 10 | 5 |
| Grid search | 10 | 5 | 10 | 5 | 100 | 30 |
| Bayesian optimizer | 10 | 4 | 10 | 5 | 100 | 30 |

**Table 3.** Spearman correlation between HyperSMURF and parSMURF scores for each of the 26 features of the Mendelian dataset.

| Feature name | HyperSMURF | parSMURF default | parSMURF grid | parSMURF BO |
|---|---|---|---|---|
| CpGobsExp | 0.13968 | 0.13819 | 0.13893 | 0.13132 |
| CpGperCpG | 0.13969 | 0.13819 | 0.13893 | 0.13132 |
| CpGperGC | 0.13968 | 0.13819 | 0.13893 | 0.13132 |
| DGVCount | 0.03413 | 0.04442 | 0.05647 | 0.06046 |
| DnaseClusteredHyp | 0.32951 | 0.32044 | 0.24895 | 0.28336 |
| DnaseClusteredScore | 0.32908 | 0.32009 | 0.24869 | 0.28315 |
| EncH3K27Ac | 0.30594 | 0.30151 | 0.24630 | 0.30958 |
| EncH3K4Me1 | 0.31055 | 0.30229 | 0.25376 | 0.31746 |
| EncH3K4Me3 | 0.23845 | 0.23367 | 0.19111 | 0.22128 |
| GCContent | 0.20083 | 0.18597 | 0.15363 | 0.19790 |
| GerpRS | 0.35444 | 0.35057 | 0.31584 | 0.30759 |
| GerpRSpv | 0.18839 | 0.18847 | 0.15510 | 0.15381 |
| ISCApath | 0.00046 | 0.00916 | −0.01227 | 0.00711 |
| commonVar | −0.18408 | −0.20035 | −0.13659 | −0.16558 |
| dbVARCount | 0.03413 | 0.04442 | 0.05647 | 0.06046 |
| fantom5Perm | 0.06291 | 0.06130 | 0.05101 | 0.05391 |
| fantom5Robust | 0.06436 | 0.06269 | 0.05217 | 0.05513 |
| fracRareCommon | 0.18258 | 0.20198 | 0.15003 | 0.17902 |
| mamPhastCons46way | 0.14808 | 0.14348 | 0.15451 | 0.16040 |
| mamPhyloP46way | 0.37928 | 0.38087 | 0.37977 | 0.43174 |
| numTFBSConserved | 0.13326 | 0.14405 | 0.14100 | 0.13788 |
| priPhastCons46way | 0.24858 | 0.25781 | 0.26809 | 0.29352 |
| priPhyloP46way | 0.33472 | 0.33576 | 0.33014 | 0.37434 |
| rareVar | 0.02916 | 0.03668 | 0.05208 | 0.05687 |
| verPhastCons46way | 0.15317 | 0.15002 | 0.16128 | 0.16888 |
| verPhyloP46way | 0.38423 | 0.38685 | 0.38594 | 0.43717 |

**Table 4.** Imbalance of the number of negative and positive examples across different regulatory region types in the Mendelian dataset. In column 4, the ratio between negative and positive samples; in bold, the lowest ratio used to under-sample the negatives for each regulatory region. The last column reports the number of down-sampled negatives for each regulatory region after applying the best ratio found in column 4.

| Category | Number of positive sampl. | Number of negative sampl. | Ratio (neg/pos) | Downsampled negative samples |
|---|---|---|---|---|
| Intronic | 75 | 5393802 | 71917 | 64625 |
| Exonic non coding | 2 | 35058 | 17529 | 1723 |
| Intergenic | 126 | 8325494 | 66075 | 108570 |
| 3' UTR | 30 | 144348 | 4811 | 25850 |
| 5' UTR | 158 | 843572 | 5339 | 136143 |
| Other | 15 | 12925 | **861** | 12925 |
| | | | **Tot. negatives:** | 349836 |

**Table 5.** Imbalance of the number of negative and positive samples across different regulatory region types in the GWAS dataset. In column 4, the ratio between negative and positive samples; in bold, the lowest ratio used to under-sample the negatives for each regulatory region. The last column reports the number of down-sampled negatives for each regulatory region after applying the best ratio found in column 4.

| Category | Number of positive sampl. | Number of negative sampl. | Ratio (neg/pos) | Downsampled negative samples |
|---|---|---|---|---|
| Intronic | 937 | 539311 | 575 | 115166 |
| Exonic non coding | 2 | 3511 | 1755 | 245 |
| Intergenic | 891 | 832571 | 934 | 109512 |
| 3' UTR | 46 | 14470 | 314 | 5653 |
| 5' UTR | 228 | 84300 | 370 | 28023 |
| Other | 11 | 1352 | **122** | 1352 |
| | | | **Tot. negatives:** | 259951 |

**Table 6.** Examples of pathogenic Mendelian single nucleotide variants where *parSMURF* sensibly outperformed *hyperSMURF*. The first two columns refer to the chromosomal coordinates of the variant. Ref and Alt to the reference and alternative allele; OMIM to the OMIM code of the associated Mendelian disease; Gene to the symbolic name of the target gene; PMID to the PubMed ID of the related publication; Region to the type of the regulatory region. The two last columns refer to the difference of ranking between respectively *parSMURF* with grid optimization and *parSMURF* with Bayesian optimization with respect to *hyperSMURF*.

| Chr | Position | Ref | Alt | OMIM | Gene | PMID | Region | Rank diff. Grid/Default | Rank diff. BO/Default |
|---|---|---|---|---|---|---|---|---|---|
| chr1 | 100661453 | T | G | MIM 248600 | DBT | 20570198 [1] | 3' UTR | 2308597 | 169786 |
| chr3 | 12421189 | A | G | MIM 604367 | PPARG | 15531525 [2] | Promoter | 663054 | 421027 |
| chrX | 138612889 | G | A | MIM 306900 | F9 | 23472758 [3] | Promoter | 194290 | 111499 |
| chr13 | 100638902 | A | G | MIM 609637 | ZIC2 | 22859937 [4] | 3' UTR | 70175 | 69069 |
| chr6 | 118869423 | A | G | MIM 609909 | PLN | 18241046 [5] | Promoter | 63078 | 55789 |
| chr16 | 31202818 | G | A | MIM 608030 | FUS | 23847048 [6] | 3' UTR | 50539 | 103623 |
| chr12 | 121416444 | T | G | MIM 600496 | HNF1A | 22413961 [7] | Promoter | 21848 | 65773 |

**Table 7.** List of newly annotated pathogenic variants used as independent test set to assess the generalization capabilities of *parSMURF*. The first two columns refer to chromosomal coordinates, REF and ALT to the reference and alternative allele (including both SNVs, micro-insertions and micro-deletions), Region the type of regulatory region, Gene the symbolic name of the gene and PMID the PubMed ID of the related publication.

| Chr | Pos | Ref | Alt | Region | Gene | PMID |
|---|---|---|---|---|---|---|
| 1 | 155261709 | G | A | exonic non-coding | PKLR | 18708292 |
| 1 | 155263324 | C | T | exonic non-coding | PKLR | 26728349 |
| 1 | 155271259 | C | G | promoter | PKLR | 18708292 |
| 1 | 155271269 | CAGAGA | C | promoter | PKLR | 26728349 |
| 3 | 10183453 | AGCGCGCACGCAGCTCCGCCCC GCGTCCGACCCGCGGATCCCGCGGC | A | 5' UTR | VHL | 30006056 |
| 3 | 10183466 | C | CTCCGCCCCGCG | 5' UTR | VHL | 30006056 |
| 3 | 38675715 | C | T | promoter | SCN5A | 27625342 |
| 4 | 89444948 | C | T | promoter | PIGY | 26293662 |
| 5 | 131705516 | G | A | 5' UTR | SLC22A5 | 31187905 |
| 6 | 100040906 | G | T | enhancer | DHS6S1 | 26507665 |
| 6 | 100040987 | G | C | enhancer | DHS6S1 | 27551809 |
| 6 | 100041040 | C | T | enhancer | DHS6S1 | 26507665 |
| 7 | 19157199 | C | T | 5' UTR | TWIST1 | 30040876 |
| 7 | 19157207 | G | T | 5' UTR | TWIST1 | 30040876 |
| 7 | 156584142 | A | T | enhancer | LMBR1 | 27592358 |
| 7 | 156585476 | C | G | enhancer | SHH | 29543231 |
| 8 | 11565816 | G | C | 5' UTR | GATA4 | 25099673 |
| 8 | 19796936 | A | G | 5' UTR | LPL | 27578109 |
| 8 | 102505149 | GA | G | 5' UTR | GRHL2 | 29499165 |
| 8 | 102505272 | CT | C | 5' UTR | GRHL2 | 29499165 |
| 8 | 102505561 | G | T | 5' UTR | GRHL2 | 29499165 |
| 9 | 21974830 | GCTCCCCGCCGCCCGCTGCCTGCTC | G | 5' UTR | CDKN2A | 26581427 |
| 9 | 21974847 | G | A | 5' UTR | CDKN2A | 26581427 |
| 9 | 21974868 | A | T | 5' UTR | CDKN2A | 26581427 |
| 9 | 21974916 | CCCT | C | 5' UTR | CDKN2A | 26581427 |
| 9 | 21975024 | GAGT | AAAG | 5' UTR | CDKN2A | 29216274 |
| 10 | 23481888 | AGCGGCGGCTGCGGCGGCGCGGCGCCG | A | exonic non-coding | PTF1A | 28663161 |
| 10 | 23508442 | A | G | enhancer | PTF1A | 28663161 |
| 11 | 31828391 | ACTT | CA | 5' UTR | PAX6 | 30291432 |
| 11 | 31828391 | AC | A | 5' UTR | PAX6 | 30291432 |
| 11 | 31828396 | C | T | 5' UTR | PAX6 | 30291432 |
| 11 | 31828461 | TAA | T | 5' UTR | PAX6 | 30291432 |
| 11 | 31828474 | CT | C | 5' UTR | PAX6 | 30291432 |
| 11 | 31832375 | C | T | 5' UTR | PAX6 | 30291432 |
| 11 | 67250359 | CCG | C | promoter | AIP | 20506337 |
| 13 | 110802675 | G | T | 3' UTR | COL4A1 | 27666438 |
| 13 | 110802678 | C | T | 3' UTR | COL4A1 | 28369186 |
| 13 | 110802678 | C | A | 3' UTR | COL4A1 | 27666438 |
| 13 | 110802679 | C | A | 3' UTR | COL4A1 | 27666438 |
| 15 | 23810779 | GTCAG | G | promoter | MKRN3 | 29763903 |
| 15 | 23810849 | C | T | promoter | MKRN3 | 30462148 |
| 15 | 35080829 | A | G | 3' UTR | ACTC1 | 27139165 |
| 17 | 10536907 | C | T | splicing | MYH3 | 31491409 |
| 17 | 10559406 | C | T | 5' UTR | MYH3 | 31491409 |
| 17 | 41277375 | T | A | 5' UTR | BRCA1 | 30075112 |
| 17 | 61996359 | G | T | promoter | GH1 | 27252485 |
| 17 | 70117348 | G | A | 5' UTR | SOX9 | 28546996 |
| 18 | 28683379 | C | G | promoter | DSC2 | 27531918 |
| 19 | 2249105 | CA | C | promoter | AMH | 31238341 |
| 19 | 49468612 | GG | CT | 5' UTR | FTL | 28636169 |
| X | 38202566 | G | T | enhancer | OTC | 29282796 |
| X | 38211793 | T | G | 5' UTR | OTC | 29282796 |
| X | 38211808 | G | A | 5' UTR | OTC | 29282796 |
| X | 38211811 | A | G | 5' UTR | OTC | 29282796 |
| X | 38211834 | C | T | 5' UTR | OTC | 29282796 |
| X | 38211835 | C | T | 5' UTR | OTC | 29282796 |
| X | 38211844 | C | A | 5' UTR | OTC | 29282796 |
| X | 70442966 | C | CT | | GJB1 | 28283593 |
| X | 70443186 | G | T | | GJB1 | 28283593 |
| X | 70444424 | C | T | 3' UTR | GJB1 | 29236290 |
| X | 85302634 | G | A | promoter | CHM | 28271586 |
| X | 85302634 | G | T | promoter | CHM | 28271586 |
| X | 138612871 | AC | A | promoter | F9 | 27865967 |
| X | 147031110 | T | C | 3' UTR | FMR1 | 26554012 |
